# Cell tracking with accurate error prediction

**DOI:** 10.1101/2024.10.11.617799

**Authors:** Max A. Betjes, Sander J. Tans, Jeroen S. van Zon

## Abstract

Cell tracking is an indispensable tool for studying development by time-lapse imaging. However, existing cell trackers cannot assign confidence to predicted tracks, which prohibits fully automated analysis without manual curation. We present a fundamental advance: an algorithm that combines neural networks with statistical physics to determine cell tracks with error probabilities for each step in the track. From these we can obtain error probabilities for any tracking feature, from cell cycles to lineage trees, that function like p-values in data interpretation. Our method greatly speeds up tracking analysis by limiting manual curation to rare low-confidence tracking steps. Importantly, it also enables fully-automated analysis by retaining only high-confidence track segments, which we demonstrate by analyzing cell cycles and differentiation events at scale, for thousands of cells in multiple intestinal organoids. Our approach brings cell dynamics-based organoid screening within reach, and enables transparent reporting of cell tracking results and associated scientific claims.

## Introduction

Cell proliferation, differentiation, movement, and their organization in complex cell lineages are key to understanding organ homeostasis and associated diseases. The development of organoid cultures, which recapitulate key features of organ development *ex vivo* [1, 2], has enabled the study of developmental dynamics at the single cell level using time lapse microscopy [3-8]. To address the complex challenge of analyzing the dynamics of hundreds of cells in dense 3D organoid architectures over multiple generations, AI-driven semi-automated algorithms have been developed that track cells based on their fluorescently labelled nuclei [3, 4, 9-12].

However, all current cell tracking approaches face a fundamental limitation: algorithms output a single tracking solution among many possible solutions, and are highly prone to making errors, yet lack a statistical basis to quantify prediction uncertainty (**figure 1A**). This lack of statistical interpretability makes rigorous analysis based on cell tracks impossible, as the inability to assess the confidence of cell tracking-based results can lead to unfounded conclusions and, more generally, limits scientific transparency and reproducibility. Finally the black-box nature of cell tracking even hampers method development and optimization itself as it makes it difficult to identify and tackle the true source of tracking errors. In contrast, other widely-used bioinformatics methods like sequence alignment [13, 14] or differential gene analysis [15], do inherently provide statistics on their output, with the resulting confidence in data interpretation and reporting crucial to their widespread adoption.

**Figure 1.**
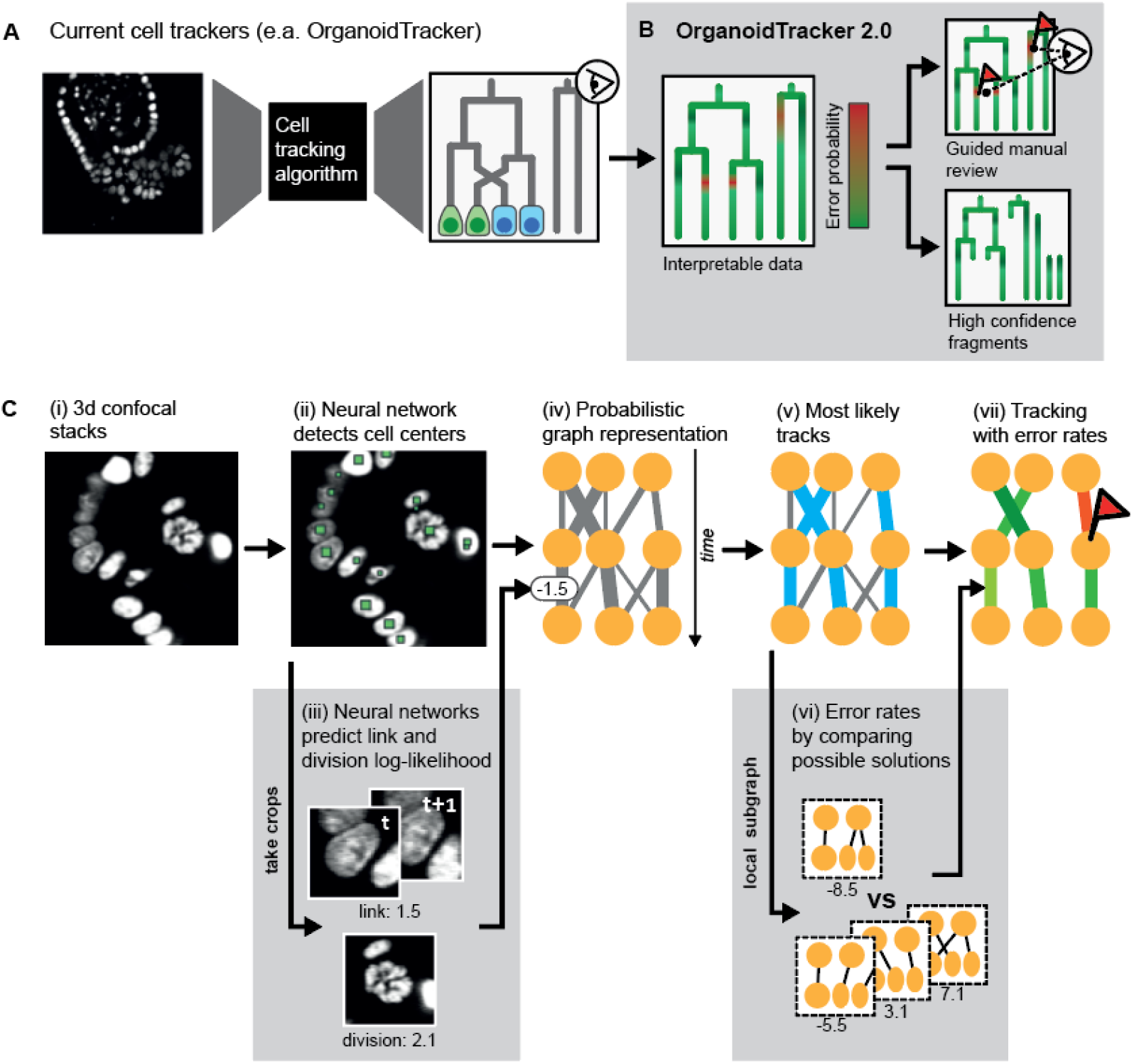
Method overview. **A)** Current cell tracking algorithms convert microscopy images into cell tracks without providing information on accuracy. Yet, even single errors can greatly alter the biological interpretation of lineages (here, change in symmetry of divisions). Hence, extensive manual review is required and finally no assessment of statistical confidence can be provided. **B)** OrganoidTracker 2.0 outputs not only tracks but also associated error rate estimates, which can function like p-values, greatly aiding data interpretability and transparency. These error estimates also enable a drastically reduced manual review, or fully automated filtering to achieve high confidence datasets. **C)** Method workflow, highlighting two novel components (grey boxes). (i) Generation of 3D confocal stacks of nuclear marker fluorescence. (ii) Neural network detection of nuclei centers. (iii) Neural network prediction of cell linking or division likelihoods, based on image crops. (iv) Constructing a graph representation of the tracking problem, based on predicted link and division likelihoods. (v) Determination of the global maximum-likelihood solution representing the most likely cell trajectories. (vi) Estimating link error rates through systematic comparison with alternative tracking solutions. (vii) Predicted cell tracks with error rate predictions for individual links.

These problems are particularly acute in context of development and tissue homeostasis, where error in even a single tracking step can radically alter the biological interpretation (**Figure 1A**). For organoids, additional tracking challenges are presented by closely-packed nuclei that move rapidly during cell division [16]. While the recent adoption of neural networks in cell tracking algorithms has greatly increased tracking quality [4, 9, 10], current methods are far from error-free, especially in organoids [3, 4]. Existing methods use *ad hoc* heuristics, like rapid nuclear volume changes or large cell displacements, to flag potential errors for manual correction [3, 4]. These methods rely on manually set cut-offs and user interpretation, which hampers reproducibility and assessment of data quality. Moreover, because such heuristics cannot provide any measure of confidence for the obtained cell tracks, creating error-free datasets relies on extensive manual curation, up to the point of checking essentially each tracking step. This process can take days for a single 300-500 cell organoid, which burdens users and limits tracking applications. In particular, screening different growth conditions or mutant backgrounds is prohibitively time consuming to analyze.

Here, we present a conceptually new approach: an algorithm that determines both cell trajectories and their error rates (**Figure 1B**). Building on our previously developed OrganoidTracker [4], we introduced two major innovations: first, we show that neural networks can perform key tracking tasks while outputting ‘calibrated’ likelihoods, meaning they provide accurate estimates of error probability of their prediction. This enables splitting the tracking procedure into specific sub-tasks that can be independently optimized, while their calibrated likelihoods can be combined to give a single, global likelihood. We developed dedicated neural networks for the sub-tasks of linking cells between subsequent microscopy images and that of identifying division events. Second, we used concepts from statistical physics, including microstates, partition functions and marginalization, to compute ‘context aware’ error probabilities. In practice, this procedure implements our intuition that a low-likelihood tracking step can in fact be of high-confidence, if all alternative cell linking arrangements are excluded by high-confidence tracks of surrounding cells.

Importantly, these innovations now also enable the reporting of statistical significance. The resulting OrganoidTracker 2.0 provides error probabilities for any lineage feature of interest, from cell cycles to entire lineage trees. These error probabilities can function like p-values in assessing the likelihood that tracking features are correct. Our innovations also enhanced tracking performance. First, OrganoidTracker 2.0 is a highly competitive cell tracker, with output tracks containing errors at <0.5%/cell/frame for intestinal organoid data, even before manual curation. Moreover, it drastically sped up this manual curation, by focusing it on those parts of cell tracks that had high predicted error rates. A 60 hour movie with over 300 cells tracked for over 300 time points was curated in ∼3 hours rather than 3 days. Second, the resulting method enables fully automated analysis without any human curation, by removing, instead of reviewing, the low-confidence parts of cell tracks and using the high-confidence parts for further analysis. Demonstrating the power of this approach, we extracted cell cycle time and proliferation rates for 20 organoids in fully automated manner, thus opening up the possibility of high-throughput screening of cellular dynamics. We show that OrganoidTracker 2.0 also provides excellent automated tracking of C. *elegans* embryos, with its performance here ranking as the best performing tracking algorithm on the Cell Tracking Challenge *[17]*. Furthermore, we provide an easy user interface, extensive documentation and straightforward retraining procedures for different biological model systems. Our framework is modular and can thus readily incorporate other cell tracking functions, such as automated cell death detection[18]. Finally, the statistical basis provided by our method will be important for data sharing and statistically justifying scientific claims.

## Results

### Method overview

Our method is divided in two parts: first, we use neural networks to identify the cells in each frame and predict the likelihoods of all possible links between them (**Figure 1C**, i-iv). Next, we use these results to find the most likely tracks and compute their associated error rates (**Figure 1C**, v-vii). Central to our approach is the concept of a probabilistic graph description of the tracking problem [19] (**Figure 2A**). Here, each node represents a cell detected at one time point, as obtained by a neural network that predicts cell center locations from 3D images of fluorescently labeled nuclei. Links between the nodes represent possible connections between these cell detections, through movement or division. To each link, we assign a ‘link energy’, defined as the negative relative log-likelihood of a link being true, so that a low energy indicates a more plausible link. Similarly, we determine a ‘division energy’ for each node that indicates the likelihood of division. A key innovation is that we employ neural networks to predict these link and division likelihoods based on microscopy data. Here, we leverage the property of neural networks that their output scores can function as accurate estimates of relative log-likelihood [20], which has thus far rarely been used in tracking applications. Using an integer flow solver, we can find the paths on the graph with the minimal associated energy [19]. This collection of paths then corresponds to the maximally likely set of cell trajectories. Finally, we use the link energies and graph structure to compute error probabilities for every link in the predicted tracks, thereby providing us with both the trajectory predictions and their associated error rates. Below, we discuss each step in more detail.

**Figure 2.**
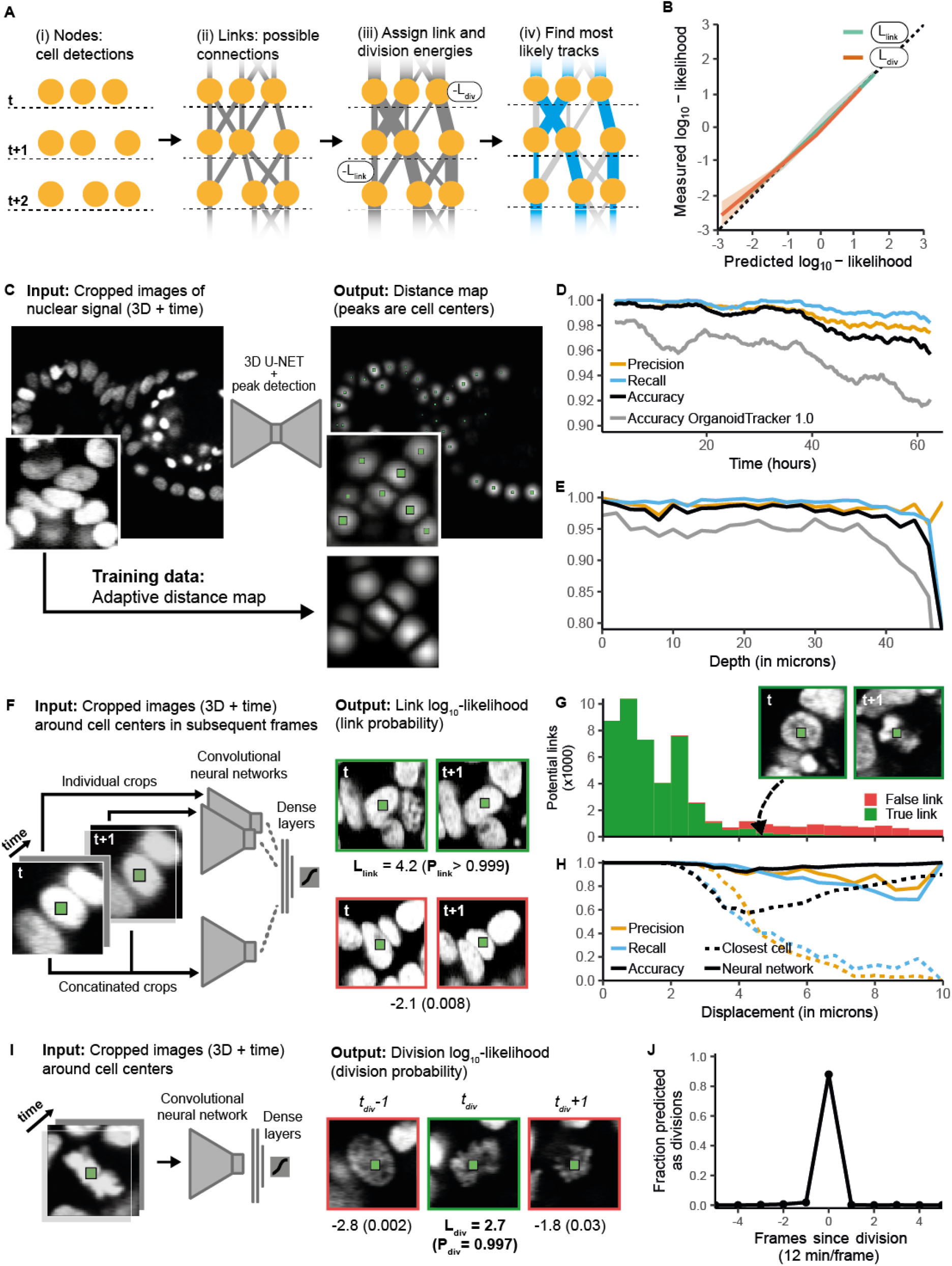
Probabilistic graph construction by neural networks. **(A)** Probabilistic graph workflow. Nodes are detected cells, and grey lines are possible links that connect cells between subsequent time points, reflecting either movement or division. Thicker lines indicate more likely links, or a lower ‘energy’. Blue lines indicate the global maximum-likelihood solution. Cell detection (i) and link and division likelihood prediction (iii) is performed by neural networks. **(B)** The predicted relative log-likelihoods by the neural networks strongly correlate with measured relative log-likelihoods (the probability of being true in the manually annotated control), both for links and divisions. The dotted line corresponds to perfect calibration. Data represents n=5 organoids, with shaded region denoting standard deviation around the mean.**(C)** 3D-UNET neural network trained to generate a distance map that indicates proximity to nuclei centers. Cell centers (green squares) are obtained by peak finding. **(C), (D)** Accuracy of cell detection, compared to OrganoidTracker 1.0, as function of (C) time and (D) imaging depth, for one organoid data set. Metrics are averaged over 10 frames. Only cells <40 um deep were included for (C). **(F)** Convolutional neural network trained to predict link likelihoods, based on crops centered around two cells detected in subsequent time points. Output images demonstrate a high (green) and low (red) likelihood link prediction, corresponding to a true and false link, respectively. **(G)** Link analysis for manually curated data. Links representing small cell displacements are typically true (green) and large-displacement links false (red). Both true and false links are observed in between, demonstrating that displacement alone does not determine link correctness. Images show a correct large-displacement link, of a cell undergoing division. (**H)** Link prediction accuracy. For all displacements, the neural network strongly outperforms predictions based on the “smallest displacement” criterion, which assigns the link that minimizes displacement as correct. **(I)** Neural network trained to predict division likelihood based on image crops centered at detected cells. Images show three subsequent frames of a dividing cell: just before chromosome separation (green border, high predicted likelihood), and before and after division (red, low predicted likelihood). **(J)** Fraction of cell crops assigned as dividing (>50% likelihood) versus time relative to division, defined as the last time point before chromosome segregation. Division assignments occur predominantly at the exact measured division time.

### Cell detection

We detect cell centers using a 3D-UNet neural network [21] (**Figure 2C**). Specifically, this network uses 3D images of organoids carrying a fluorescent nuclear marker (H2B-mCherry, **Supplementary figure 1**) to predict a distance map, which for every pixel records its distance to the closest cell center [4, 10]. Cell centers are then obtained by finding local peaks in this distance map. This approach enables the generation of training data by annotating cell centers, which is less labor-intensive than manual 3D segmentation of nuclei. An inherent challenge to distance maps is that cell center peaks for closely-packed nuclei blend into one another, causing under-segmentation. We therefore developed an adaptive distance map, in which we assign increased distance values to pixels that are almost equidistant to two cell centers (**Figure 2B, Supplementary figure 2A**). This ensured that cells remain well-separated in the resulting map, thus reducing segmentation errors (**Supplementary figure 2B**). To test the impact of the adaptive distance map on cell center detection (together with additional improvements in the training data generation and augmentation pipeline, see **Methods**), we compared performance to OrganoidTracker 1.0. Both methods were trained on the same data set and subsequently tested on identical unfamiliar microscopy images. Overall, detection accuracy was improved substantially, decreasing the error rate ∼4-fold over the complete time-lapse data set. The high accuracy only decreased slightly, from 99% to 95%, when the cell’s nuclear fluorescence had poor signal-to-noise ratio, for example after prolonged imaging (>50 hours, **Figure 2D**) or deep in the imaging volume (>40 μm, **Figure 2E**).

### Estimating link and division likelihood

We then construct the linking graph by connecting each node, representing a detected cell, with all possible links. For practical purposes, we culled links representing unrealistically large displacements (**Methods**). Links either connect the same cell in two consecutive frames, or connect a mother and daughter cell. The first category of cells only move and generally retain similar nuclear shape and fluorescence intensity between frames, while dividing cells show distinct changes in nuclear morphology. To deal with this heterogeneous data, we trained a neural network to estimate the relative log-likelihood of a link being correct, based directly on the imaging data. Here, we employ a fundamental ability of classification neural networks that use a cross-entropy loss during training, especially when combined with Platt scaling subsequently [20, 22], namely that the probability they predict exactly matches the actual probability of the event being correct as observed in ground-truth data (typically referred to as a ‘well-calibrated likelihood’). This property of neural networks has thus far not been used in cell tracking and is key for our error rate estimation.

We designed a neural network that inputs cropped 3D images centered on each cell’s detected position for time points *t* and *t+1*, and predicts the likelihood that they represent the same cell, using information within the crop on both the nucleus and its local environment (**Figure 2F**). The trained network correctly assigned low energy (high likelihood) to links between the same cell, even when the fluorescence signal changed substantially, while assigning high energy (low likelihood) for links connecting a cell to its neighbor. We compared the network’s performance to the baseline criterion, often used for tracking [23], that the links representing the smallest displacement between frames are correct. First, when comparing all possible links in the graph against ground-truth data, we found that links corresponding to small displacements (<3 μm) were often correct (**Figure 2G**), while large displacement links (>7 μm) were incorrect. However, many correct links represented substantially larger displacement (3-7 μm). These large displacements occurred predominantly during cell division (**Figure 2G**, inset, **Supplementary figure 3**), and their correct assignment is thus essential for lineage tree reconstruction. We therefore compared the accuracy of links assigned by the neural network (link considered true when predicted likelihood>50%) to those assigned by the ‘smallest displacement’ criterion (for each cell, only the shortest-distance link towards it is true) as function of link displacement. For <3μm displacement, both approaches yielded similar high accuracy (**Figure 2H**). However, the accuracy decreased strongly for larger displacements when using the ‘smallest displacement’ criterion, but remained high for the neural network, also for cells close to division (**Supplementary figure 3**), demonstrating the superiority of our neural network approach.

**Figure 3.**
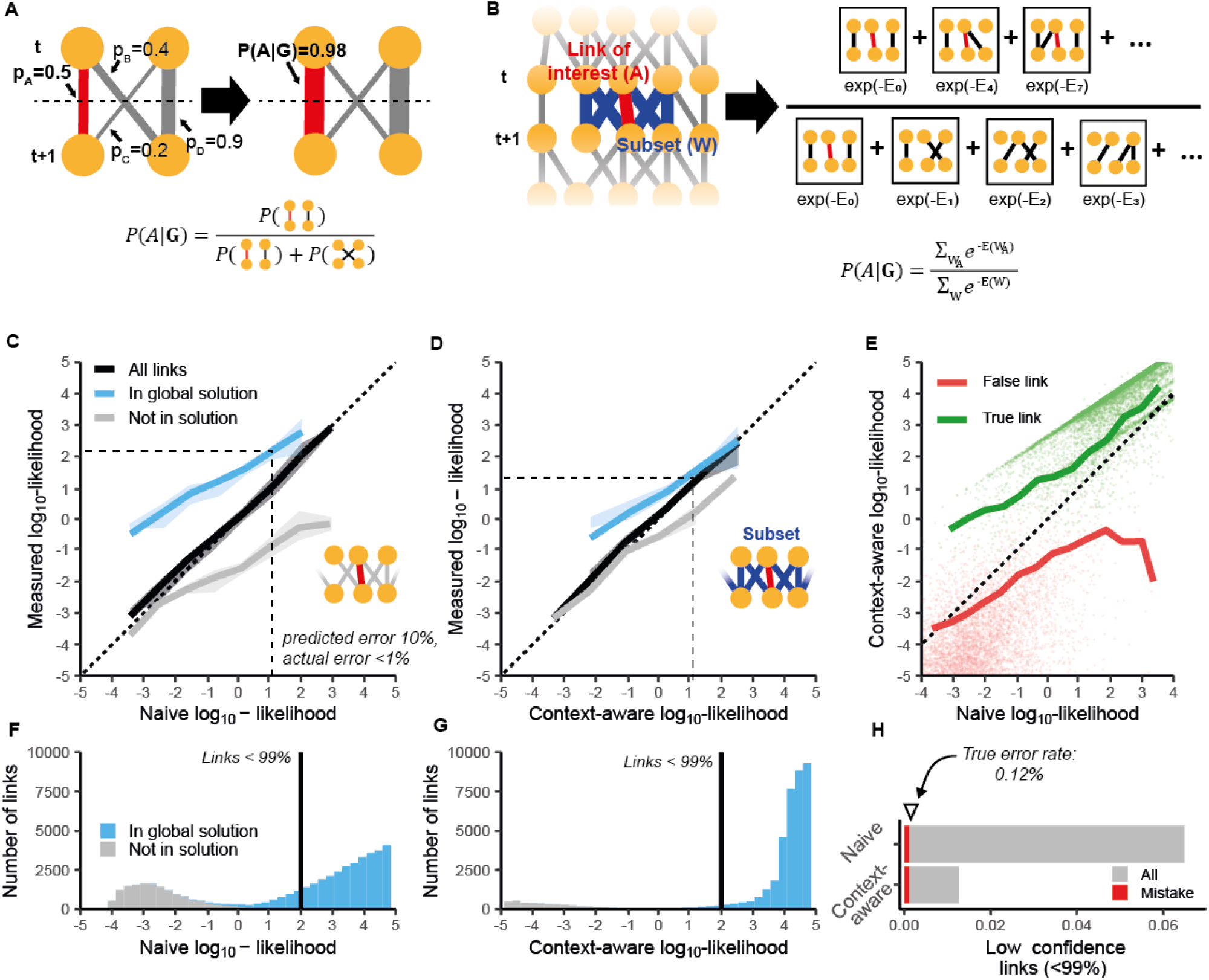
Error rate estimation by marginalization. **(A)** Simplified example explaining marginalization. Lines indicate putative links *A*-*D*, with thickness indicating their likelihood. Link *A* (red) has low predicted likelihood. However, as links *B* and *C* are unlikely because *D* has high likelihood, it implies *A* is true with high certainty. The probability P(A|G) that *A* is true given graph structure *G*, can be calculated by looking at the probability of configurations containing link A versus ones that do not. (**B)** Schematic outline of marginalization, performed on a subset of links around the link of interest. P(A|G) is given by the summed energy of all configurations containing link *A* normalized by the summed energy of all configurations. **(C)**,**(D)** Measured link likelihood versus (C) naive likelihoods predicted by the neural network or (D) context-aware likelihoods calculated by marginalization. Data is shown for all possible links (black) or links that are either in the global maximum-likelihood solution (blue) or not (grey). For naive likelihoods (C), links in the tracking solution are more likely correct than expected on the basis of the neural network prediction, while for context-aware likelihoods (D) they more closely match measured likelihoods, reflecting integration of graph information. Dotted line is perfect calibration. Data for n=5 organoids. Shaded region is S.D.M. (**E)** Context-aware likelihoods versus naive likelihoods. Dots are individual links and lines are averages, for true (green) or false links (red). Marginalization increased predicted likelihood of correct links while decreasing it for incorrect links. **(F)**,**(G)** Number of links versus predicted (F) naive or (G) context-aware link likelihood. While for (F) most links in the maximum-likelihood solution (blue) are predicted as high-confidence (>99% likelihood), a fraction have confidence levels similar to rejected links (grey). In contrast, for (G) virtually all maximum-likelihood links are now predicted as high-confidence. (**H)** Fraction of links in the maximum-likelihood solution deemed low-confidence (<99% likelihood) after marginalization. Red is the fraction of low-confidence links that were actual errors compared to ground-truth, while the arrow indicates the fraction of errors in all links. Almost all errors that occurred were covered by the <99% likelihood threshold. Marginalization thus reclassified many low-confidence links as high-confidence, but not those that represent errors.

While most graph nodes have a single correct outgoing link, reflecting movement, nodes of dividing cells have two correct links connecting to its two daughters. Dividing cells exhibit distinct nuclear morphology, with chromosomes forming the metaphase plate, which we exploited to determine the likelihood that a node corresponds to a dividing cell and thus allowed to have two outgoing links. We designed an additional neural network that used 3D image crops to predict division likelihood, including the previous and subsequent frame to facilitate identifying the exact moment of division (**Figure 2I**). When testing images at different times relative to division, defined as the last frame before chromosome separation, division assignment (>50% likelihood) indeed coincided with the moment of division in >90% of cases (**Figure 2J**). Moreover, cells at time points before or after division were only rarely assigned as dividing, even though they are visually similar to cells at the exact time of division.

We also substantially improved prediction by up-sampling challenging cases during training: fast-moving, dividing cells for link prediction (**Supplementary figure 4**) and time points around division and cell death for division prediction (**Supplementary figure 5**), as dying cells resembled dividing cells in appearance. This general ability to tailor training data sets to individual tasks (cell detection, link and division detection) is a major advantage of our modular approach, compared to merging multiple tasks in a single, more complex neural network [3, 10]. This fine-tuning also shows the benefits of having directly interpretable probabilities, instead of generic weights[4, 10], as the output of tracking subroutines in guiding method optimization. Finally, we validated that neural network output indeed represented true probabilities. We binned all possible links based on their predicted likelihood to be correct. Then for each bin, we calculated the true likelihood, i.e. the fraction of links that were correct according to the ground-truth data. We found that predicted likelihoods were well-calibrated, with predicted and true link likelihood matching for the full likelihood range (**Figure 2B**). This ability to accurately predict likelihoods is essential for estimating tracking errors, as discussed below.

**Figure 4.**
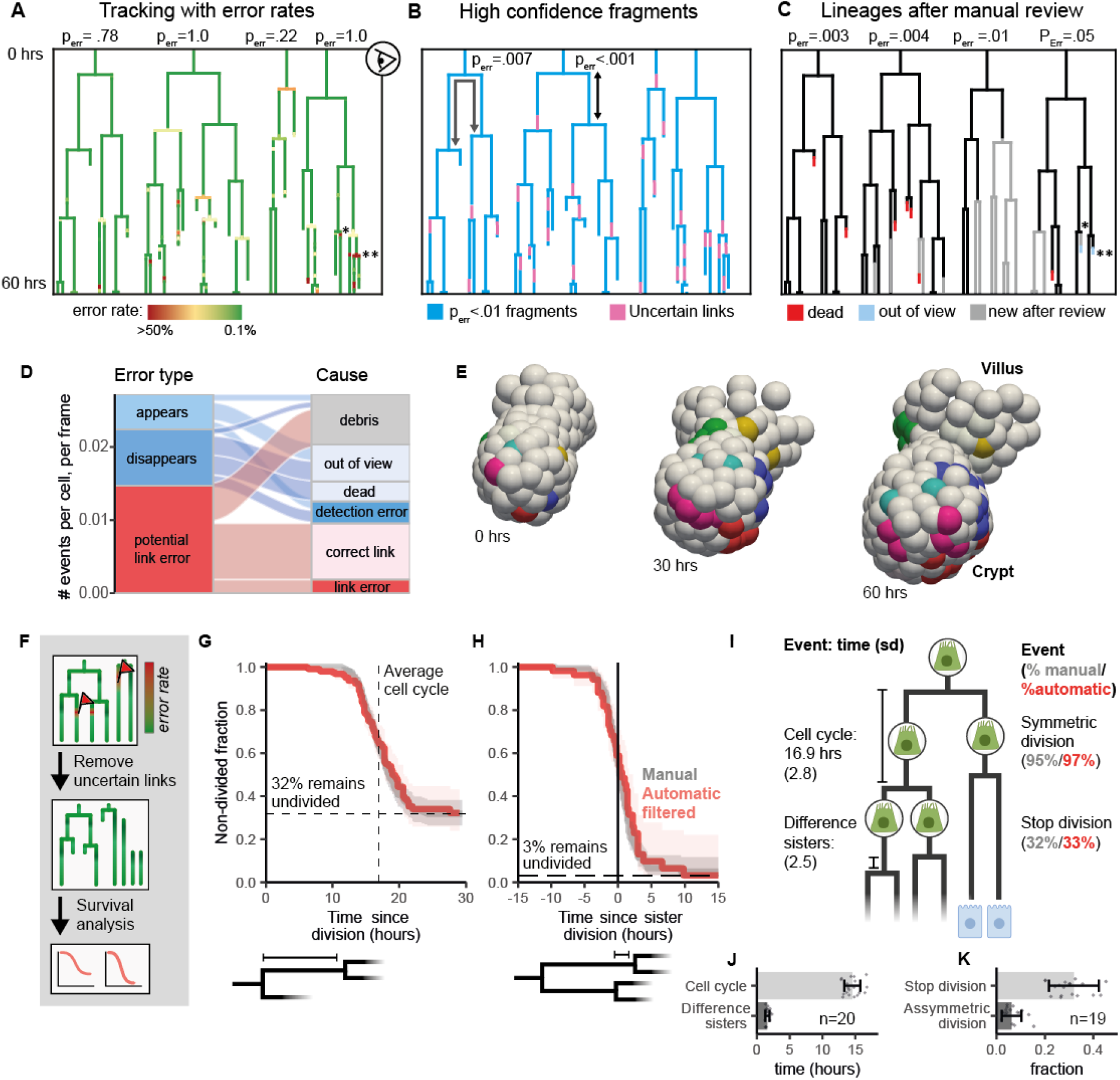
Applications. **(A)** Selected lineages before review, with color indicating associated error rates. Low-confidence links are yellow to red, (**) indicates a flagged error associated with a clearly erroneous lineage structure (unrealistically short cell cycle), while (*) would not be identifiable as an error based on lineage structure alone. Error probabilities are calculated for each lineage tree by combining error rates of all underlying links, identifying lineage trees as low-confidence even though they visually appear plausible. (**B)** High-confidence _l_ineage fragments (<0.01 error rate), obtained based on individual link error rates. Users can identify high-confidence cell cycles (black arrow) or dividing cells that are sisters (grey arrow) without manual review. **(C)** Lineage trees after manual review of potential errors flagged in (A). Grey lines: lineage sections added as consequence of curating erroneous links. P-values now indicate high confidence that lineages truly reflect underlying cell dynamics. **(D)** Characterization of potential errors. Links flagged as potential errors either represent (dis)appearing cells (blue) or are low-confidence (red). A significant proportion of potential errors represented very short tracks of cellular debris, whose correction impacted the data only minimally. Only a minority of potential errors are actual errors in cell detection or linking that required correction. **(E)** 3D reconstructions including lineages in (C). Colors indicate cells in the same lineage. **(F)** Automated analysis without manual review, by filtering out low-confidence links and performing survival analysis on the resulting, partly censored data. **(G)** Survival curve indicating the probability that a cell has not divided at time *t* after its birth. Shown are manually annotated (grey) and automatically filtered data (red) for all cells in a single organoid. Vertical line denotes average cell cycle duration, while horizontal line shows inferred fraction of cells that stop dividing after birth. Shaded region is 95% confidence interval. **(H)** Survival function indicating the probability that a cell has not divided, as a function of the time relative to the division of its sister. Almost all cells divide within a 10 hour window around the time of sister division, showing that symmetric divisions dominate intestinal organoid growth. Shaded region: 95% confidence interval. **(I)** Comparison of key lineage dynamics parameters obtained by fully automated (red, no manual review) or manual analysis (grey). Fully automated analysis shows excellent agreement with manually annotated data. **(J)** Automatically obtained cell cycle times (grey) and differences between sisters (dark grey) for twenty organoids. **(H)** Automatically obtained fractions of cells that stop dividing, for all cells (grey) and cells that have a dividing sister (dark grey) for nineteen organoids.

### Maximum-likelihood track prediction

To construct cell trajectories, we use a min cost flow solver algorithm [19] to select the set of links in the probabilistic graph that globally maximize likelihood (minimize energy). While close to optimal tracks are obtained readily, the algorithm does not guarantee identification of the global optimum and we typically found minor mistakes, such as link pairs that increase global likelihood when swapped. Moreover, flow solvers cannot change the graph structure by adding or merging nodes, causing vulnerability to under- and over-segmentation[24]. We therefore automatically check if overall likelihood is improved by swapping link pairs and by adding or removing nodes (**Methods**). In the latter case, we identify pairs of disconnected tracks that can be joined by adding a node, corresponding to a missed cell detection, or by merging nodes, corresponding to over-segmentation (**Supplementary Figure 6A**,**B**). New links and nodes are assigned energies reflecting the probability of under- and over-segmentation, as given by the detection network’s recall and precision. Overall, these additional steps substantially increase the duration over which cells can be continuously tracked and show the benefits of our probabilistic framework in data post-processing (**Supplementary figure 6C**).

### Context-aware estimation of link error

Central to our approach is estimating the error rate of individual links. The ‘naive’ link likelihoods, as predicted by the neural network, in principle provide information on each link’s error probability, but ignore the context of the link likelihoods predicted for surrounding cells. The importance of context is evident already in manual tracking: here, human trackers typically first establish high-confidence links, which in turn, by reducing the remaining possible links, facilitates subsequent assignment of lower-confidence links. Such contextual information can be computed in our probabilistic framework as well. This is illustrated by the simplified graph in **Figure 3A**, where the likelihood of a low-confidence individual link is increased dramatically (from 50% to 98%) in the context of the larger graph, because high-likelihood links exclude all alternative linking arrangements. To generalize this notion, we considered the total likelihood of all possible linking arrangements containing the link of interest, using the following approach (**Figure 3B**): for each possible linking arrangement, corresponding to a tracking solution *W*_*i*_, we calculate its energy *E*(*W*_*i*_) by summing all link and division energies, with the solution likelihood proportional to 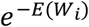, The ‘context-aware’ likelihood link *A* is then given by the total likelihood of all tracking solutions containing that link, 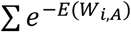, normalized by the sum for all possible solutions, 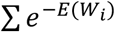 We call this procedure marginalization, as all other variables are marginalized out to arrive at a single-link error rate estimate without referencing other any links. Since summing over all possible tracking solutions is computationally unfeasible, we restrict this to local subgraphs, here consisting of links are less than 3 steps away (**Figure 3B, Methods**), thereby balancing performance and computational complexity, with marginalization requiring <1 hour for a 60 hour time-lapse data set.

The above approach borrows conceptually from statistical physics, with each possible tracking solution equivalent to a microstate and the normalization factor to the partition function. From a probabilistic perspective, our method resembles the multiplicative opinion pooling framework [25, 26], where different opinions (here, neural network predictions) are combined assuming that their pooled likelihood is proportional to the product of likelihoods associated with the individual opinions. Here, we have extended this framework to predictions that concern different events (link and divisions) that are part of a larger structure (the graph), instead of simply reflecting the same event (see **Methods & Supplementary text)**.

Our marginalization procedure assumes that the individual link and division likelihood predictions are independent. This is not strictly true, as predictions are based on shared information: for example, the various image crops used as input can overlap. This causes overconfident error predictions (**Supplementary figure 7**). As a correction, we employ the similarity with statistical physics to introduce a ‘temperature’ *T* that decreases energies to *E*_*i*_ /*T*, thereby reducing confidence levels. These scaled energies represent the unique information contained in each prediction. The correct scaling factor is obtained phenomenologically as the one for which the marginalized error rate predictions are well-calibrated, based on the same data sets used for neural network training and validation (**Supplementary figure 7, Methods**).

### Evaluation of error rate predictions

We compared both naive and context-aware error rate predictions against measured error rates, by testing them against manually annotated data sets. To avoid bias, these datasets were generated completely independently from the OrganoidTracker pipeline. We then reused the ground truth cell centers to generate the link predictions and calculated the context aware error rates. First, we examined naive link predictions, i.e. before marginalization. Overall, naïve predicted likelihoods were well-calibrated, correlating strongly with measured likelihood (**Figure 3C**). However, our data indeed indicated that the error predictions are under-confident in determining true and false links without access to contextual tracking information. Specifically, links identified by the flow-solver as part of the global maximum-likelihood solution displayed measured likelihoods much higher than naive neural network predictions, while for links rejected from the global solution measured likelihoods were lower than predicted (**Figure 3C**). This matches our intuition that the graph contains additional information on link likelihood (**Figure 3A**), as the flow-solver selects links based on the complete graph information, while the neural network makes its naive predictions based only on local image information. In contrast, context-aware link predictions showed strongly improved confidence, reducing the mismatch between predicted and measured likelihood for both flow-solver selected links and the rejected links (**Figure 3D**). Moreover, incorporating graph context specifically increased the predicted likelihood of true links, while decreasing it for false links (**Figure 3E**).

The improved context-aware link predictions have significant practical advantages for error correction. For naive predictions, a large fraction (6%) of links selected by the flow-solver must be reviewed, when using <99% predicted likelihood as threshold for manual curation. At the same time there was significant overlap in likelihood values between the flow-solver selected links and those it rejected (**Figure 3F**), erroneously suggesting that these links were equally likely to be true. However, for context-aware predictions, the predicted likelihoods of flow-solver selected links differed strongly from rejected links (**Figure 3G**). Moreover, the number of <99% confidence links that must be manually reviewed reduced substantially (1%), while practically all true linking mistakes (0.12%) were still detected (**figure 3H**). An order of magnitude more links (∼25%) must be reviewed to achieve similar accuracy using a simple heuristic based on cell displacement (**Supplementary figure 8**).

As a final control we tested the marginalization procedure in the context of our full pipeline. We ran our algorithm on three different organoids. We then performed manual curation to generate ground truth data to which the predicted error rates could be compared. Again we found that the error rates were well-calibrated (**Supplementary figure 9)**. Lastly, we could reduce the computation time without reducing accuracy by excluding highly unlikely links before marginalization, as this did not significantly impact likelihood prediction and error rate computation (**Supplementary figure 9, Methods**).

### P-values for lineage features and manual curation

We ran our pipeline on a ∼60 hour time-lapse dataset of a representative organoid. Marginalization yielded cell tracks and lineages with an error rate for each link (**Figure 4A**). The rate of potential errors (defined as <99% likelihood links) was 1.5%/cell/frame, after removing tracks deep in the imaging volume, with potential errors predominantly, but not exclusively, concerning divisions (**Figure 4A)**. These error rates can now be used to compute error probabilities for complex lineage features as *p*=1 ∏_*i*_ *P*_*i*_, where *P*_*i*_ are the context-aware likelihoods of all links *i* of the lineage feature of interest, with *p* representing the probability that the feature is not correctly tracked (**Supplementary figure 10)**. These error probabilities enable users to assess the statistical significance of, for example, individual cell cycles, the observation that cells are sisters, or even entire lineage trees (**Figure 4A,B, Methods**), functioning similarly to p-values. Moreover, we identified high-confidence lineage fragments as fragments with *p*<0.01 calculated over all links. We found that long stretches of unprocessed tracking data was of such high-confidence, with high-confidence fragments often spanning multiple cell cycles (**Figure 4B, Supplementary figure 11**).

Our accurate knowledge of link likelihoods implied that minimal manual correction focusing only on low-confidence links should be sufficient to obtain error-free tracks. To test this approach, we manually reviewed all links with <99% likelihood. We also reviewed all beginnings and endings of cell tracks mid-experiment (0.9%/cell/frame), which represented either cell death, cells entering or exiting the field of view or, potentially, cell detection errors. Of these potential errors, only a fraction represented true linking or detection errors (0.3%/cell/frame, **Figure 4D**). However, correcting the few true errors did strongly improve the lineage trees, complementing them with previously unconnected sub-trees. (**Figure 4C, Supplementary figure 12**), underscoring the importance of identifying even infrequent tracking errors when complete lineage trees must be produced. Finally, independent manual tracking of cells in the lineages of **Figure 4C** yielded identical trees. Correction required ∼4 hours for a data set where ∼300 distinct cells were tracked in a ∼60 hour time window (**Figure 4E, Supplementary figure 12**). When we calculated error probabilities for cell lineages after manual correction, assigning a probability of one to manually corrected links and recomputing the marginalized link-likelihoods (**Methods)**, we generally found low values, P<0.01 for all analyzed lineage trees (**Figure 4C**), indicating high resulting confidence.

### Fully automated lineage tracking by error filtering

Current 3D cell tracking algorithms require extensive manual curation after automated cell tracking before analysis is performed. Our ability to accurately estimate linking error rates enables a novel and fundamentally different approach: to remove low confidence track fragments and subsequently perform analysis only on the remaining high-confidence fragments, without any human oversight (**Figure 4B and F**). For fragments that have high-confidence data from division to division, properties such as cell cycle durations can be directly measured and compared between different organoids and regions within an organoid (**Figure 4B**). We also employed survival analysis (**Methods**), a statistical framework for dealing with censored data [27-29], to quantify a broad range of lineage properties while also incorporating information from lineage fragments containing incomplete cell cycles. Specifically, we generated Kaplan-Meier survival curves to estimate the fraction of non-divided cells as a function of time since cell birth (**Figure 4G**), using all (incomplete) high-confidence track fragments that included at least one birth. This survival curve displayed two notable features: first, it plateaued at 32%, representing the fraction of cells that do not divide again and, hence, have differentiated. Second, the average cell cycle time is given by the time at which 50% of the dividing fraction have divided, with standard deviation corresponding to the spread around this time, yielding a cell cycle duration of 17±2.8 hour. We then extended this analysis to sisters, using sister pair fragments to generate survival curves as a function of time relative to sister cell division (**Figure 4H**). Here, the curve plateaued at 3%, representing the fraction of sisters where one proliferated while the other ceased proliferation. This high degree of symmetry between sister cells is consistent with recent work [5, 6]. Moreover, sisters had highly similar cell cycle durations, as indicated by the steep decline in the curve, corresponding to a difference in cell cycle duration of 2.5 hour between sisters. Overall, survival curves generated from automatically filtered and manually tracked data showed almost exact overlap (**Figure 4H**). Finally, we demonstrated the automated nature of this approach by analyzing 20 different organoids (**Figure 4J and K**). We consistently found similar parameter values and survival curves, even as organoids displayed notable differences in size and morphology, indicating that the underlying lineage dynamics is independent of this morphological variation (**Figure 4J and K, Supplementary figure 13**).

### Out-of-sample capabilities

While our methodology was designed and optimized for intestinal organoids, we also examined the performance on other samples, specifically a confocal time-lapse microscopy dataset of *C. elegans* embryogenesis hosted by the *Cell Tracking Challenge* [17, 30]. This was chosen as it was the only available confocal microscopy dataset of a 3D embryo. We trained cell detection, division, and link prediction neural networks with only minimal changes to training procedure (see **Methods**). We found that our method here performed as well as for intestinal organoid data, generating cell tracks spanning up to 7 generations (**Supplementary figure 14)**, even though training data was limited in comparison. Upon manual review of all <99% confidence links and (dis)appearance events, corresponding to 0.9% of total links, the resulting data exactly reproduced the overall *C. elegans* lineage structure. Importantly, we also verified that our method produced well-calibrated error rates, by comparing against them against the manually corrected data. Independent verification of our automated tracking results, before any correction, by the *Cell Tracking Challenge* confirmed the quality of our predictions, ranking us first in tracking performance out of 18 competitors. Our TRA-score (a metric of tracking performance with a maximum of 1[31]) of 0.995 represents a more than two-fold improvement over the closest competitors[9, 10]. Overall, this demonstrates that our approach is readily applied to other multicellular systems.

## Discussion

In this study we presented a conceptual innovation in cell tracking: whereas existing algorithms typically generate tracks with minimal information on correctness, OrganoidTracker 2.0 instead estimates the confidence in its predictions. Our approach exploits neural networks to predict linking and division likelihoods based on 3D microscopy data, and uses a probabilistic graph representation of the tracking problem to adjust these likelihoods based on information of surrounding cells. This also enables the computing of error probabilities, functionally equivalent to p-values, for any tracking feature, thereby reporting their statistical significance. This enables both highly efficient manual curation, by only correcting a minority of low-confidence tracking steps, as well as fully automated analysis, by using only high-confidence track fragments. In this way, we quantified key lineage parameters without human supervision and at high throughput for multiple organoids. The ability to compute error probabilities will be important to improving scientific transparency and reproducibility. Our method allows researchers to report the statistical significance of the cell tracking results that they produce, and the associated scientific claims, which is of increasing relevance in science. We believe these capabilities will be important in further stimulating the adoption of cell tracking methods, and the quantification of cell dynamics within tissue systems. Our approach is readily extended to cell tracking in other contexts, such as 2D cultures or embryos. OrganoidTracker 2.0 is freely available, with extensive documentation and an user-friendly GUI [4].

Our approach uses neural networks not only for cell detection, but also to estimate the probability that cells are linked or will divide between two time points. Here, the key insight is that these link and division neural networks can be trained to predict well-calibrated likelihoods, meaning that the predicted probability of a putative link or division being correct matched the probability of it being part of the manually annotated ground truth (**Figures 2**,**3**). Predicting likelihoods necessitated a modular design, with separate neural networks for detection, linking and identifying divisions, rather than a single network for detection and linking simultaneously [3, 10]. This modularity brings further advantages. First, this enabled task-specific optimization both of network architecture and training data, for instance by up sampling the number of challenging division events when training the division network. This optimization is greatly aided by the fact that these subtasks have easily interpretable probabilities as their output. Second, each network can be swapped with other implementations [32-37] tailored to different model systems, as long as they provide well-calibrated probabilities. Finally, it allows extending our approach with additional neural networks, to predict likelihood of other events that impact cell tracking, such as cell death, cell extrusion or abnormal divisions [18].

Our ability to predict error probabilities represents a fundamental advance in the cell tracking field. Current state-of-the-art 3D cell tracking typically rely on heuristic rules to identify tracking errors, such as flagging unrealistically large displacements or short cell cycle times [3, 4]. Recent 3D cell tracking algorithms used neural networks for cell linking [36] and detection [35] that provided approximate information on link and division likelihood, but not in way that supports calculating error rates and statistical significance. For 2D cell tracking, studies used approaches such as linear regression, Bayesian analysis, random forests or Kalman filters [32-34, 38-40] to predict link and division likelihoods, sometimes even explicitly calibrating these outputs [33], but do not provide error rates or otherwise quantify statistical significance based on these. The key enabling step here is our marginalization procedure (**Figure 3**), which incorporates the intuitive understanding that likelihoods must be adjusted based on tracking information from surrounding cells. For instance, a low-likelihood link can still be of high confidence if all possible other linking possibilities are ruled out by high-likelihood links to other cells. Without marginalization, link likelihood is typically under-confident, resulting in too many links erroneously ranked as low-confidence for the error probabilities to be useful for subsequent analysis (**Figure 3**). Our marginalization procedure is independent of how link likelihoods are calculated and hence can be readily incorporated in other (cell) tracking algorithms.

The current best-practice approach to address inevitable cell tracking errors is to follow automated tracking by labor-intensive manual review [3, 41]. Importantly, our accurate error rate prediction strongly reduced the time required for manual correction, by focusing review exclusively on the few links with >1% error rate. Consequently, a 60 hour time-lapse movie of intestinal organoids with ∼300 cells, required only 4 hours of manual review (**Figure 4**), while this would take several days otherwise. Our predicted error probabilities also enable fundamentally novel analysis and experiments, without any human supervision: by focusing exclusively on high-confidence cell tracks or lineages, one can fully automatically extract lineage features and relationships. Using this approach, we extracted key features of cell proliferation control, such as cell cycle length, cell cycle arrest rate and cell cycle correlations between sister cells, in a fully automated manner and at high throughput (20 organoids, with a computation time of ∼1 hour per organoid on a desktop computer). Automated analysis of high-confidence cell track fragments has many other possible uses. First, it can be extended to other biological events, such as cell death or cell cycle stages, when combined with fluorescent markers [29, 42] or neural networks that can detect it [18]. Moreover, it enables systematic characterization of cell proliferation parameters or other features under different conditions [43], such as the addition of signaling inhibitors or drugs. Finally, it can also be used to study processes such as cell movement and rearrangement, by quantifying for example the rate at which adjacent cells separate over time[6].

Our results raise fundamental issues regarding the reporting of cell tracking-based results. For small data sets, manual curation may be performed at least on a limited number of key features such as divisions. However, for larger data sets this approach is no longer feasible, such as for larger embryo or gastruloid systems [44], or when comparing organoids across many different conditions or genetic backgrounds. Yet, once established, reported tracking results are often treated as a given – without insight into the uncertainties. Currently the only way for readers to assess the confidence of these results and associated claims is to study the original microscopy images and perform (manual) tracking oneself, which is typically not feasible. The ability to calculate error probabilities equivalent to p-values, as we advance here, will be of general importance to mitigate this issue. Like any other form of quantification in science, such error probabilities or error probability cut-offs should be reported for displayed cell tracks, lineage trees, and for lineage features, such as cell cycles. Reporting error probabilities of published tracking data will also be crucial for data sharing, by enabling external users to assess confidence in different features of the data, even without access to the underlying microscopy images. Our work here now provides the conceptual framework and computational tools to extend this approach to a broad range of cell tracking applications.

Many salient features of aberrant organ development and homeostasis, including cancer, are only, or most readily, visible on the level of single-cell dynamics. Examples include (local) cell proliferation and death rates, spatial heterogeneity in proliferative potential, and how these are impacted by drugs and mutations. While recent studies demonstrate microscopy-based screens of cancer organoid shape and size [45], single-cell analysis at scale is not yet feasible [46, 47]. Our demonstration of fully automated cell-tracking analysis is therefore of direct relevance to cancer research, and opens a new frontier in understanding tissue homeostasis in disease.

## Methods

### Organoid culture

Mouse intestinal organoids with a H2B-mCherry reporter were used, gifted by Norman Sachs and Joep Beumer (Group of Hans Clevers, Hubrecht Institute). Organoids were grown embedded in membrane extract (BME, Trevingen) in medium consisting of murine recombinant epidermal growth factor (EGF 50ng/ml, Life Technologies), murine recombinant Noggin (100ng/ml, Peprotech), human recombinant R-spondin 1 (500ng/ml, Peprotech), n-Acetylcysteine (1mM, Sigma-Aldrich), N2 supplement (1x, Life Technologies) and B27 supplement (1x, Life Technologies), Glutamax (2mM, Life Technologies), HEPES (10mM, Life Technologies), Penicilin/Streptomycin (100U/ml 100μg/ml, Life Technologies) in Advanced DMEM/F-12 (Life Technologies). Organoids were kept in incubators at 37°C and with 5% CO2. The medium was changed every two days. Each week organoids were mechanically broken and the fragments reseeded.

### Sample preparation

Organoids were seeded around two days before imaging in 4 well chambered cover glass (#1.5 high performance cover glass) from Cellvis. In order for the organoids to locate within the lens working distance and minimize the required laser power, we placed the sample on a cold block (∼4°C) for ten minutes after seeding. In this way the organoid fragments could sink to the bottom before the gel solidified. Afterwards the BME gel was allowed to solidify at 37°C for 20 minutes before adding medium.

### Microscopy

Imaging was performed on a Nikon A1R MP microscope with a 40x oil immersion objective (NA = 1.30). Around 30 z-slices with a 2 µm step size were taken per organoid every 12 minutes, with a pixel size of 0.32 µm^2^.

### Training data

Our training data consisted of 9 different tracked crypts together with nearby villus regions. Timelapses were between 16 hours and 65 hours long and with the full dataset totaling 281 hours (1405 frames). This is the same dataset used to train the original OrganoidTracker [4] so we can confidently say that any improvements are due to the new algorithm and not because of an expanded training dataset. All training data was generated in the context, and partially used in an earlier publication [5].

### General neural network training and prediction procedure

The input during both training and predicting for all neural networks consists of a list, in which each item references an image frame together with any data needed to create the final neural network input (i.e. a list of cell centers around which to crop). Only during training and prediction image frames are loaded and is the input data generated in order to minimize memory footprint. All data augmentation during training is done at run-time for the same reason. Image frames can be loaded from .tiff files but also from common platform specific file formats like .*lif* (Leica) or .*nd2* (Nikon) to avoid the need for data conversion.

Before training the neural network, the input list is randomized and split into a training and validation set (80% vs. 20%). After training the link and division detection data we perform a simple Platt scaling based on the validation dataset to make sure our predictions are well calibrated [22].

Gradient descent is done using the ADAM-optimizer for all neural networks. The full network architectures can be found on our GitHub (https://github.com/jvzonlab/OrganoidTracker).

### Cell center detection – generating training data

To detect cell centers we use both the frame at the time point of interest and the subsequent frame to give the neural network access to dynamic information. We crop the images to a box which contains all annotated cell centers to avoid learning on unannotated regions. Images are then normalized after which random crops (32 × 96 × 96 × 2t) are made. Users can set arbitrary time windows and crop sizes when training their own neural networks.

To augment the data these crops are randomly flipped along the x or y-axis (50% of cases) or randomly rotated and scaled (by a random factor between 0.8 and 1.2). Further augmentation is done by randomly changing the contrast by exponentiation of the intensity values by a random number (between 0.8 and 1.2). The fluorescence intensity decay with increasing image depth can vary greatly between imaging settings. We therefore also augment the data by increasing the decay in intensity with depth by a random factor, such that the deepest frame can have up to 4-fold reduction in intensity

### Cell center detection – distance map and weights

The neural network is trained to predict for every pixel in the image the distance to the nearest cell center. The distances are transformed by a Gaussian function, to give rise to diffuse spots centered around cell centers. This approach has achieved success in many cell localization algorithms where the full segmentation of cells is not available [4, 10, 48]. We improve this approach by also taking into account distances to nearby cells other than the closest one. By increasing the distances (and thereby decreasing intensities in the distance map) for pixels that are close to another cell, we ensure that the Gaussian spots remain well separated. The mathematical description of the ‘adaptive’ distance *d* is given by:

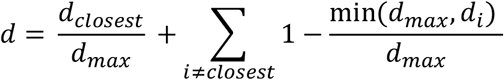

In which *d*_*max*_ represents the maximum radius within cell centers are still relevant in computing the distance value for a pixel. It can be chosen up to the minimum distance between two cell centers before spots will overlap. The first term measures the distance to the closest cell center, while the second term increases this value if other cell centers are also within *d*_*max*_.

The intensity values in the distance map are then given by:

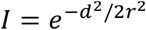

In which we choose *r* to be 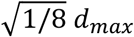 to produce well separated spots.

The calculation of the distance map is done at run-time on the GPU for maximum efficiency. It can be implemented using only convolutional operations by replacing the minimum operator in the equation above by a pseudo-minimum (soft-min function).

Our algorithm allows users to only partially annotate datasets reflecting the fact that most existing manually tracked data is often focused on a limited region of interest due to time considerations. The training on partial annotations enabled by assigning large weights to pixels in the annotated regions versus the background during training. To assign these weights we change our distance map so that pixels with multiple cells nearby have lower distance values associated with them:

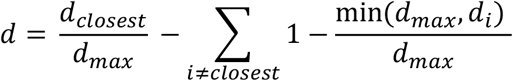

We then use these distances to calculate the weight values:

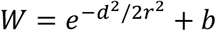

where *b* is a small weight assigned to background pixels. By giving some weight to the background the neural network can learn to ignore debris and imaging artefacts outside of the foreground. For the intestinal organoid data *b* is chosen such that half of the total summed weights is associated with annotated nuclei and half with the much larger background region.

### Cell center detection – neural network

The neural network used for cell detection is very similar to the 3D-UNET used in the previous OrganoidTracker [4]. The different time-points in the input are treated as different channels. A new element in the network is a final smoothing layer (convolution with Gaussian kernel with a 1.5 pixel width). Because the center point annotation is inherently noisy (not pixel perfect) the predicted output should be smooth. By enforcing this explicitly we reduce overfitting and speed up the training.

### Cell center detection – peak finding

From the predicted distance map, we localize the cell centers by using a peak finding algorithm, as described before [4]. Peaks within a certain radius (half the typical distance between nuclei) of other higher peaks are excluded by the peak finding process to avoid oversegmentation due to noise in the predicted distance map.

During cell division cells round up and their distance to other nuclei increases. At the same time cells are more prone to oversegmentation as the H2B fluorescence is not uniformly distributed anymore because of chromosome condensation. To counteract this, we revisit the cell detections after we have predicted the division probabilities (see below) and merge dividing cell detections (defined as having a division probability greater than 50%) that are closer than 5 µm from each other.

### Cell center detection – Evaluation

Cell center detection was evaluated as previously done [4]. We compared predicted data with partially annotated manual datasets. The evaluation data consisted of five different organoids, imaged on different days, for which at least one crypt was fully tracked. The organoids were tracked for between 90 and 320 frames.

For every cell center in the manually annotated dataset we check if there is a predicted cell center within 5 µm, these count as true positives. A predicted center can only match a single cell center in the manual data. Unmatched manual annotations are false negatives. Predicted cell centers that remain unmatched and are within the manually annotated region (5 µm distance from an annotation) are counted as false positives. Consigning the evaluation to annotated regions means that mistakes far from the epithelial layer are ignored (i.e. debris recognized as nucleus), but these are both rare and generally irrelevant for tracking.

Recall is calculated by dividing true positives by the total number of manual annotations. Precision is defined by dividing the false negatives over the amount of predicted cell centers within the annotated region. Accuracy is the number of mistakes over the sum of all observations (true positives, false positives, false negatives).

To test the effect of our ‘adaptive’ distance map we also trained a network on a target mapping that consisted simply of Gaussian spots around the cell centers. In order for these spots to not overlap we had to half their radius relatively to the ‘adaptive’ version. The pixel weights were kept the same (**Supplementary figure 2**).

### Link detection – proposing possible links

To avoid looking at extremely implausible links we propose links based on the distance between the subsequent cell detections. During both training and prediction, we only consider links from a cell detection to a cell detection in the next frame that are at most two times further away in distance than the closest cell in the next frame.

### Link detection – generating training data

The input of the neural network for link prediction consists of a crop centered around a cell center, a crop around the cell detection in the subsequent frame and a vector describing the distance in pixels between the cells. The two crops are 16×64×64 in size and both contain the two time points containing the cell center detections. Users can set arbitrary time windows and crop sizes when training their own neural networks.

Data is augmented in the same way as during cell center detection, except that we do not vary the decay in intensity with depth as the crops are much smaller in the z-dimension. Instead we increase the range in which we vary contrast (exponentiation by a number between 0.5 and 1.5).

To aid prediction we provide the neural network with direct information about the direction of movement by adding the displacement vector to the neural network inputs, besides the crops around the cell centers. It is known that convolutional neural networks have trouble integrating information in the form of Cartesian coordinates [49]. We therefore add an extra three channels to both crops. These contain for each pixel the x, y and z distance respectively to the other cell center detection in the proposed link.

We upsample difficult cases, cells that are dividing (within a window of an hour around cell division) or move a considerable distance (more than 3 µm less than 7), by a factor of 5.

### Link detection – neural network

The first part of the neural network for link detection consists of two convolutional neural networks (CNNs). To maximize the amount of information extracted one CNN takes in the concatenated crops while the second CNN takes as input a single crop (two identical copies of the second CNN are available to analyze both crops). This means that one CNN can integrate pixel information between crops and directly assess how alike the two cell detections at subsequent time points are. The other CNN is forced to focus on a single crop, which could in combination with information about the direction of movement already be enough to assess the link likelihood.

The features extracted by the CNNs in combination with the displacement vector are then fed into multiple densely connected neural network layers to yield a prediction.

### Link detection – Evaluation

To evaluate the link neural network, we used the same set of evaluation data as used in evaluating the cell center detection. See main text for the evaluation procedure.

To test the effect of adapting the training data we also trained a link detection neural network without upsampling difficult cases (see ‘Link detection – generating training data’). We then compared accuracy, precision and recall across all evaluation organoids (**Supplementary figure 4**).

### Division detection – generating training data

The input of the division detection neural network is a crop (12×64×64) centered around a cell center, with the previous and subsequent frames included for dynamic information. Data augmentation is done the same as during link detection training.

To avoid a too low frequency of images related to cell division, we upsample cells in the process of division (within a one hour window around the nucleus dividing) by a factor of 10. We also upsample all dying cells (cells with tracks ending before the end of the experiment), as these can closely resemble dividing cells. From all other cell detections, which are often trivial to predict as non-dividing, only a random subset is included so that they make up 20% of the total dataset.

### Division detection – neural network

The design of the division detection neural network mimics that of the link detection network. A convolutional neural network extracts features that are then fed into a dense layer to generate the prediction. The main difference is that due to limited nature of the division datasets (there are only hundreds of divisions present in our training data) we employ only a single dense layer to avoid overfitting.

### Division detection – Evaluation

To evaluate the division neural network, we again used the same set of evaluation data used in evaluating the other neural networks. See main text for the evaluation procedure.

To test the effect of adapting the training data we also trained a division detection neural network without upsampling difficult non-dividing cases (see ‘Division detection – generating training data’). We replaced these difficult cases by randomly selected cell centers, so that divisions make up the same fraction of the training data as in our normal training procedure. If we would truly train on an unbiased sampling the data, so that non-divisions make up the vast majority, this would cause the training procedure to not converge. We then compared accuracy, precision and recall across all evaluation organoids (**Supplementary figure 5**).

### Graph description

In our graph description of the dataset we follow the framework developed by Haubold et al. [19]. Here the nodes of the graph are the detected cell centers and the edges the proposed links. These edges have an associated energy penalty that is the negative log-likelihood that the link is true as predicted by the neural network. The nodes have an associated division penalty which is again the negative predicted log-likelihood.

Within this framework we also have to assign energy penalties to the events where a track disappears or appears or that a cell detection is a false positive. A track can disappear when a cell dies or its next position is not detected. The disappearance probability is thus the combination of the death rate and the false negative rate of the neural network. Here, the latter makes the dominant contribution. Tracks can appear when their previous position is not detected, which again relates to the false negative rate. The probability of a cell detection being spurious is given by the false positive rates. All these rates can be estimated from the validation of the cell detection neural network and are around 1%. Varying these probabilities within an order of magnitude (3% to 0.03%) does not significantly affect the track prediction or the marginalization procedure (not shown).

To account for cells appearing or disappearing because they are close to edge of the imaging volume and can leave the imaging volume, we assign lower (dis)appearance penalties (corresponding to a 10% chance of (dis)appearance) to cell detections at the edges of the volume.

One could imagine a neural network that would assign explicit probabilities to the correctness of cell detections so that we could use node specific (dis)appearance penalties. This should lead to minor improvements in track quality, but such an approach would have several drawbacks. First of all, training data is limited because the cell detection network makes few mistakes. Furthermore, such a neural network would have to be retrained every time a new cell detection network is trained as it is specific to the type of mistakes that that network makes. Integrating a neural network to identify dying cells and adapt the disappearance probabilities accordingly would be more feasible [18], but of limited use due to the rare nature of cell death in our system.

In principle the predictions made by division and link detection neural networks are probabilities conditional on the correctness of the underlying cell detections, because only correct cell detections are in the training data. It is possible to assign energy penalties in such a way that they represent probabilities of a link or division conditional on the existence of the node it is coming from, by combining the chance that a link is incorrect and that its source node does not exist in a single energy penalty. This could avoid including some oversegmentations that persist over multiple subsequent frames and have high likelihood links between them in the tracking solution. But including correct links between oversegmented cells is in our case actually the preferred behavior. Not including these links would hamper our approach of solving these oversegmentations during post-processing (see below). This does mean that after marginalization we also have to interpret the predicted error rates as the chance that the two different cells associated with the detections are not linked, not the chance that the link is ‘incorrect’ because one of the two detection is due to an oversegmentation. Because oversegmentations on its own already introduce errors per definition, as a track caused by oversegmentation both has to appear out of nowhere and disappear again, this will not cause any missed errors.

### Flow solver

We use the flow solver developed by Haubold et al. [19] to find the most likely set of tracks. In order to help it converge to an optimal solution we prune the graph of high energy edges. We do this by comparing every edge to its alternatives; links having the same source or target nodes. If a link with a much lower penalty is available (>4.0 difference, corresponding with a 10,000 times more likely link) we remove the edge. This was not done during the marginalization evaluation (figure 3), where link removal like this would introduce a bias in the non-marginalized probabilities for very unlikely links. Potential divisions that have a probability below 0.01 are also removed.

The flow-solver has sometimes trouble converging or halts prematurely especially in the presence of a large number of low certainty predictions. To circumvent this, users can also use the Viterbi-style algorithm proposed by Magnusson et al.[50] as implemented by Haubold et al.

### Fine-tuning flow solver solution

Because the flow solver does not guarantee an optimal solution, we fine-tune our solution by checking for every link if removing it and replacing it with an appearance and a disappearance would lower the total energy. We then also look at pairs of links in the solution that connect two nodes at timepoint *t* with two nodes at timepoint *t+1* and check if they should be replaced with a pair of edges that connects the nodes the other way around. We perform 3 cycles of this pruning and swapping of links.

### Solving over and undersegmentation

Our probabilistic description allows us to add and merge nodes in the graph in a statistically rigorous manner to tackle the track fragmentation caused by over and undersegmentation.

Tracks that are split up due to oversegmentation are easy to diagnose in the tracking solution. The tracks should partially overlap in time (minimum of one and maximum of three frames) and should have relatively high likelihood edges that are not part of the tracking solution between them. This is, because if the tracks represent the same cell, edges between nodes in the two tracks should be likely. If the combined likelihood of an edge connecting the two tracks and the likelihood of a false positive cell detection (oversegmentation) is higher than the likelihood of a track disappearing and another appearing, we favor connecting the tracks and prune the overlapping cell detections (**Supplementary figure 6A**).

Fundamentally this solves a drawback in the graph framework for tracking where every cell detection is treated as independent evidence for the existence of a cell. If for instance a cell is oversegmented in multiple subsequent timepoints this is treated as very strong evidence that there are actually two cells present. It is obvious that this actually confers little more evidence than a single oversegmentation because these predictions are and should be highly correlated between frames. Revisiting the tracking solution allows us to treat multiple subsequent oversegmentations as a single false positive event.

Tracks split because a cell was missed are found by looking at disconnected tracks with a single frame gap between them. If their start and endpoints are within a reasonable radius (less than the typical distance between nuclei) we propose a new node that connects the tracks. This node is assigned a probability of being correct that is equal to the false negative probability. New edges in the graph representation are made to all nearby nodes with an energy penalty representing a uniform link probability (**Supplementary figure 6B**).

This correction method solves another fundamental problem with using flow solvers for tracking; they can ignore cell detections but cannot add nodes for missed cell detections. A priori it is difficult to determine where ‘helper’ nodes might need to be added and allowing cell ‘merging’ to deal with undersegmentation [19] makes the tracking problem much less constrained. We instead solve it with an easily understandable and straightforward post-processing step. Earlier cell tracking solutions have employed conceptually similar methods but have to rely on manually picked parameters to regulate post-processing in the absence of a probabilistic description [24]. Here, we use our probabilistic graph description to rigorously identify the proper post-processing steps with minimal need for user set parameters.

### Marginalization

Marginalization is done on a subset of the graph to make it computationally tractable. We assume that the most informative edges (and their associated nodes) are between the same time points as the link of interest and are the ones closest to it in space. Distance is measured by how many steps on the graph have to be made to traverse edge-wise from the target node of the link of interest (**Supplementary figure 7B**). Taking three steps as our cut-off for inclusion in the subset leads to around one hour of computation time for the marginalization procedure for over an imaging experiment of over 300 frames with over a hundred cell detections per frame.

For every node in the subset all edges that point to non-members of the subset are combined in a single edge that accounts for the total probability to connect to a node outside of the set.

After subset selection we construct a set of microstates, test which microstates fit the graph constraints and calculate their associated energy. To avoid having to check the full set of binary combinations of events (∼2^*N*^) we construct the set by varying for every target node in the *t+1* timepoint which node in the previous timepoint *t* is connected and combining all these variants. In this way the number of constructed microstates scales as 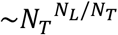, in which *N*_*T*_ is the number of target nodes, and *N*_*L*_ the number of edges (the target node can on average contact *N*_*L*_ /*N*_*T*_ possible source nodes).

Microstates can be encoded as a vector with as its length the number of events (1 if an event, like a link or division, is part of it, 0 if not). To check if a microstate is possible, we can construct a matrix that encodes the flow constraints on the graph. This matrix gives the net flow into every node when multiplied with a microstate vector. An outgoing link or disappearance event represents a flow of -1 while an incoming link or appearance gives a flow of 1. Divisions are represented with a -1 flow as they should allow an extra outgoing link. When for one or more of the nodes the flow is unbalanced, the microstate is rejected and excluded from the partition function. Total energies are calculated by taking the inner product with a vector containing the energy penalty per event. These energies are then divided by the ‘temperature’ for proper calibration (see next section).

The probability of the link of interest being true is found by dividing the sum of exponent of the negative energy of the microstates containing that link by the sum of all microstates, which is the partition function of the subset:

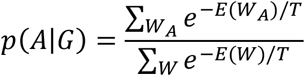

To reduce the computational burden, links that are deemed almost certainly correct (>99.99%) or incorrect (<0.01%) are marginalized over a minimal subgraph containing only the other input edges of the target node of the link in question.

### Motivation for using ‘temperature scaling’

The marginalization procedure without temperature scaling assumes that the energy penalties are derived from information that is unique to the predictor, a neural network in our case. This is not a realistic assumption as predictions might be made on the basis of overlapping crops and on shared baseline estimates. Not accounting for this overlapping information leads to overconfidence (**Supplemental figure 7**).

In our solution for this problem, we split all energies in a component that is based on shared information and in one based on information unique to that prediction. We assume that the amount of shared information is proportional to the total information about an event (*E*_*i*_):

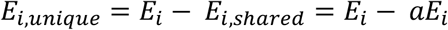

With *a* a constant between zero and one. When calculating the energy of a microstate 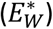 we can then sum the unique energies while assuming we can combine the shared information in a weighted manner. This weighing factor *b* (smaller then 1) should be low if all the shared information is shared between all events and higher if the overlap is less (for instance when a prediction made about a link mostly shares information with adjacent links, but not with all elements in the subset):

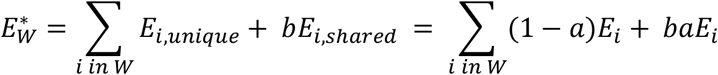

From this we derive that we can account for shared information by using a single factor that functions as a temperature (*T*). This temperature is high if much of the information in any given is not unique (high *a*) and if this shared information is shared with all other predictions (low *b*):

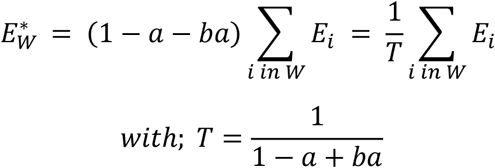

### Marginalization as an opinion pooling procedure

We can also motivate our marginalization procedure without relying on analogies with statistical physics. Instead, we can interpret our method as an extension of the ‘multiplicative opinion pooling’ framework as proposed by Dietrich [25, 26]. The idea of combining predictions in a machine learning context has an older history (Hinton, 1999), but the specific framework of Dietrich and List enables us to neatly deal with prior probabilities and overlapping information. This will prove to be key in producing well-calibrated outputs.

Multiplicative opinion pooling suggests that opinions of different agents (different predictions by neural networks in our case) can be combined by multiplying them:

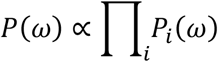

In which *ω* is a state in the set of possible states Ω and *P*_*i*_ denotes the probabilities predicted by individual predictors. Shared information between predictors can be incorporated in this framework by normalizing the predictions by the predictors’ priors based on the shared information. Conceptually this means that predictors first arrive at a consensus *P*_0_ on the basis of their shared prior information after which their unique information is pooled multiplicatively.

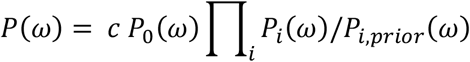

With *c* functioning as a normalization factor:

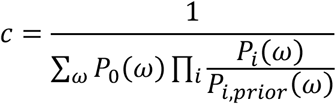

In this framework each predictor must have an opinion on all possible states. In our case predictors make only a single prediction on an event *a* (a link or division) that is part of a state. Therefore, we redefine multiplicative pooling as:

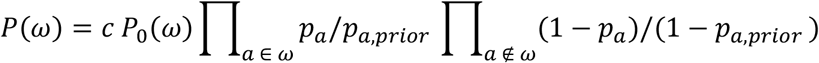

In which the microstate probability is now proportional to the product of the probabilities that its constitutive parts are true and the other events are false. The probability of a given event can then simply be calculated as:

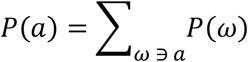

By extending multiplicative pooling in this way we retain a major motivation behind multiplicative pooling, namely ‘individualwise bayesianity’. This axiom states that is should not matter to the final prediction if extra information is integrated before or after the pooling procedure, as the input information is the same. In our case this holds on two levels (see supplementary text for the proof). Firstly, it does not matter when we introduce information about a microstate when calculating its probability (*P*(*ω*)). It also does not matter when information about an individual event is introduced when we are calculating its probability (*P*(*a*)). This provides large flexibility in post-hoc integration of new opinions, such as the judgement of a human reviewer.

The question remains how to extend our concept of ‘temperature’ to this framework. For simplification we can rewrite everything in terms of likelihoods:

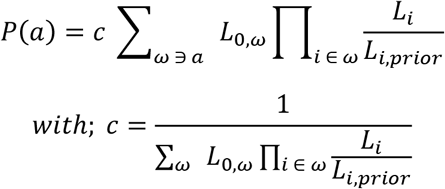

The question now remains how to define the consensus prior *L*_0_ and determine the priors. Dietrich and Listz suggest using a geometric mean on the priors if the shared information is completely shared between all agents [26]. In our case this is not necessarily true so we let the weight associated to a single prediction be free (*b*) instead of 1/*n*. For the priors we again assume that the shared information is proportional (with a factor *a*) to the total information held by an agent.

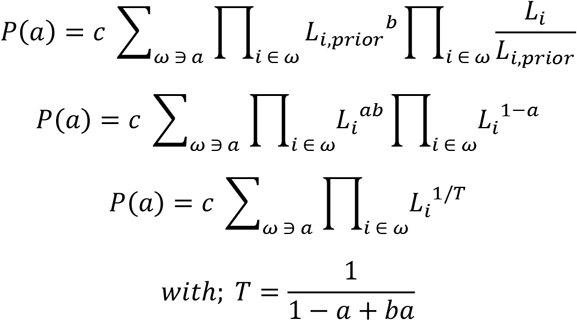

Which is equivalent to the description we arrived at using the statistical physics framework.

We finally want to contrast this opinion pooling procedure with updating a ‘Bayesian belief matrix’, a (cell) tracking approach that uses link probability estimates to connect non-dividing object detections [32, 51]. This method cannot integrate division probabilities and can only take one type of constraint into account; the fact that cells cannot merge. In situations where these are the only constraints present (for instance when considering a subgraph where only one cell is present in the later timepoint) we show that this approach is equivalent to our marginalization method (see supplementary text).

### Estimating the calibration temperature

We find the optimal temperature (as defined by the binary cross entropy loss) by calibrating on the training data. To do this we use the neural networks to predict link and division probabilities for the cell detections in the training data. Then we perform the marginalization and compare the marginalized link probabilities to the manual tracking. The task is now to find a ‘temperature’ (*T*), for which the predictions *p*_*i*_ are closest to the ground truth (*l*_*i*_ denotes the truth value of a link):

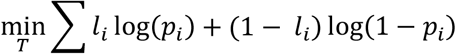

With *p*_*i*_ given by:

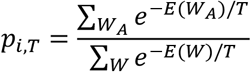

In practice most of the energy contribution in our marginalization comes from a handful, often two microstates. As an example, the dominant microstates of an uncertain link often take the form of an option where all cells move half a cell to the left and another where they move half a cell to the right.

For a given link *A* one of these states dominates the microstates that contain the link (*W*_*A*_) and the other the ones that do not contain it (*W*_\*A*_). This allows us approximate the marginalized probability *p*_*i,T*_ as:

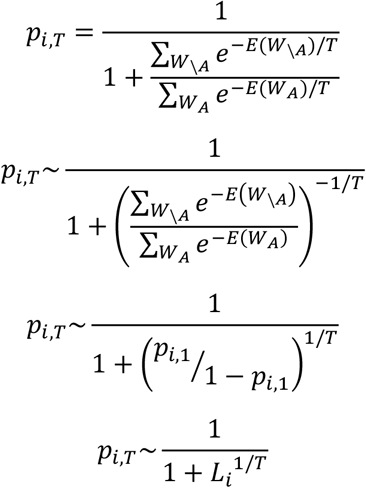

This clearly and conveniently maps on a linear regression problem, where we have to learn parameter 1/*T* given the original marginalized relative likelihoods (*L*_*i*_) as an input.

The temperature obtained this way works well on data that is part of (**supplementary figure 7**) and outside of (see next section) the training dataset in producing well-calibrated error rates. This proves that the simplifications made to arrive at a single correction factor amenable *T* to linear regression are allowable.

### Evaluation of marginalization procedure

To evaluate the correctness of the marginalized error rates we again compare our predictions against the five fully manually annotated organoids used for the other evaluations. We use the manually annotated cell centers as the input for our division and link detection and perform marginalization afterwards. This allows us to compare all error rate estimates to a fully human derived ground truth, without the need to map machine predicted cell centers on the cell centers annotated by humans. These mappings are not trivial and errors in these mappings can strongly skew the results. Furthermore, because the human assigned links are completely independent from the algorithm output, we deem this the strongest test for our marginalization procedure.

We bin the marginalized link prediction in groups based on their relative log-likelihood (15 bins). For every bin we then compute the average probability of the link being correct and compare this to actual amount on correct link in this bin as determined by looking at the ground truth.

We also perform this evaluation on the manually reviewed tracking data (see next section). Here we started out with cell centers predicted by a neural network. Verifying that the error rates are well-calibrated in this case shows that the marginalization procedure is not dependent on human annotated cell centers. We compare the error rates against tracking data where all links are corrected, but no under- or oversegmentations are solved (**Supplementary figure 9**). Solving segmentation errors involves changing the graph representation and thus introduces links without an associated error rate prediction, making evaluation impossible.

### Manual review

In order to evaluate manual annotation, we reviewed the possible errors for 3 complete organoids tracked for around 100 to 300 frames. Potential errors were flagged at all links that had a marginalized probability below 99% and the start and endpoint of appearing and disappearing tracks respectively. We first corrected all potential link errors and used the corrected data to check the calibration of the marginalized predictions as described above. We then checked all other errors and identified their cause for the largest (>300 frame long) dataset.

Error correction was done in our GUI [4], that zooms in on errors and tells user about the kind of error they encountered; possible link mistake, track appearing or track disappearing.

### *C. elegans* cell tracking

We obtained the *C. elegans* embryo datasets from the cell tracking challenge website [17]. We trained new cell detection and link and division prediction neural networks on the two provided annotated training datasets. As all cells in the imaging volume were annotated there was no need to crop the image during cell detection training and we could increase the background weighting to 0.95 without risking training on unannotated cell centers. Due to the difference in nucleus sizes compared to the intestinal organoid data we also increased the radius parameters in the distance mapping for the cell detection. All other networks were trained as for the intestinal organoids. We estimated the proper scaling temperature for the marginalization by calibrating on the training data as described for the intestinal organoid data.

We used one of the unannotated ‘challenge’ datasets to evaluate tracking quality and validate that the marginalized probabilities were well-calibrated. We did this by manually checking all potential errors and using the corrected dataset as our reference.

### Automated lineage dynamics analysis

For the automated analysis we first filtered out all links that had a marginalized probability below 99%. All tracks that do not end in a division are considered censured.

A key assumption underlying survival analysis is that the probability of an event happening is independent of the chance of being lost to follow up. In our case this assumption is broken as cells are relatively often lost when they are close to dividing (due to rapid nucleus movement) and cell division is the key event when studying lineage dynamics. This means we would underestimate the number of dividing cells, because we tend to lose track of them just before they divide. We break this dependency by using a division detection neural network to check for every track that is lost to follow-up if it is lost during the division process. We then reassign tracks which end in a predicted division (>50% predicted probability) from the censured category to the divided class. Now observing the division events is no longer affected by uncertainties in the tracking during the division process.

The neural network trained for this task was trained in the exact same way as described before except that we are not interested in pinpointing the exact moment of chromosome separation. We want also want to classify tracks as dividing if they are lost during any other moment of the division process. We therefore classify all cells within two frames around division as dividing during training. Because of the varying length of the division process, we exclude time points directly around this window to avoid including cells in the training data that look clearly mitotic but are just outside the window.

We can also use this neural network to split tracks that contain a division, but where in initial tracking no division was assigned due to lack of a plausible daughter cell, for instance because one of the daughters moves out of view. Therefore, we break up tracks when the chance of division is on average higher than 99% for three consecutive frames.

Our method detects some cells with very short cell cycles, where the cell division generally leads to cell death and not in two daughter pairs, potentially reflecting polyploid cells. These are not classified as dividing in the manually annotated data so we remove these very short cell cycles (below 6 hours). This has the added benefit that it also removes some cases where the division neural network wrongly assigns a division to a track end. Although the chance of that happening is low; it happens generally in less than 2,5% of tracks.

For survival analysis we use the ‘surv’ package in R and for the fitting ‘survflexcure’. During analysis we only use tracks that start in a division and use the next division as the event under study. Cell cycle times are analyzed by fitting a Guassian hazard to the data allowing for a ‘cured’ fraction, that will not divide again. The mean of the Gaussian represents the average cell cycle and its standard deviation the spread around this mean. The ‘cured’ fraction is used as an estimate of the fraction of differentiating cells. Prior to fitting we remove outliers that are more than ∼7 hours from the mean (more than 3 times the standard deviation). To avoid dealing with negative times we fitted a log-normal distribution to the exponents of the survival times instead of using a normal distribution directly.

Manual data for comparison is analyzed in the same way, but the only censoring events derive from cell death, the end of the experiment or cells leaving the imaging volume. No neural network thus has to be used to check if censured tracks end in a division.

## Software availability

The organoidtracker software is freely downloadable from GitHub (https://github.com/jvzonlab/OrganoidTracker). The models used are available on Zenodo (DOI: 10.5281/zenodo.13912685).

## References

1. Betjes, M.A., et al., Cell Tracking for Organoids: Lessons From Developmental Biology. Frontiers in Cell and Developmental Biology, 2021. 9.

2. Beumer, J. and H. Clevers, Cell fate specification and differentiation in the adult mammalian intestine. Nature Reviews Molecular Cell Biology, 2021. 22(1): p. 39–53.

3. de Medeiros, G., et al., Multiscale light-sheet organoid imaging framework. Nature Communications, 2022. 13(1): p. 4864.

4. Kok, R.N.U., et al., OrganoidTracker: Efficient cell tracking using machine learning and manual error correction. PLOS ONE, 2020. 15(10): p. e0240802.

5. Huelsz-Prince, G., et al., Mother cells control daughter cell proliferation in intestinal organoids to minimize proliferation fluctuations. eLife, 2022. 11: p. e80682.

6. Zheng, X., et al., Organoid cell fate dynamics in space and time. Science Advances, 2023. 9(33): p. eadd6480.

7. He, Z., et al., Lineage recording in human cerebral organoids. Nature Methods, 2022. 19(1): p. 90–99.

8. Ender, P., et al., Spatiotemporal control of ERK pulse frequency coordinates fate decisions during mammary acinar morphogenesis. Dev Cell, 2022. 57(18): p. 2153-2167.e6.

9. Sugawara, K., Ç. Çevrim, and M. Averof, Tracking cell lineages in 3D by incremental deep learning. eLife, 2022. 11: p. e69380.

10. Malin-Mayor, C., et al., Automated reconstruction of whole-embryo cell lineages by learning from sparse annotations. Nature Biotechnology, 2023. 41(1): p. 44–49.

11. Wait, E., et al., Visualization and correction of automated segmentation, tracking and lineaging from 5-D stem cell image sequences. BMC Bioinformatics, 2014. 15(1): p. 328.

12. Bragantini, J., M. Lange, and L.A. Royer, Large-Scale Multi-Hypotheses Cell Tracking Using Ultrametric Contours Maps. ArXiv, 2023. abs/2308.04526.

13. Mitrophanov, A.Y. and M. Borodovsky, Statistical significance in biological sequence analysis. Briefings in Bioinformatics, 2006. 7(1): p. 2–24.

14. Altschul, S.F., et al., Gapped BLAST and PSI-BLAST: a new generation of protein database search programs. Nucleic Acids Res, 1997. 25(17): p. 3389–402.

15. Soneson, C. and M. Delorenzi, A comparison of methods for differential expression analysis of RNA-seq data. BMC Bioinformatics, 2013. 14: p. 91.

16. McKinley, K.L., et al., Cellular aspect ratio and cell division mechanics underlie the patterning of cell progeny in diverse mammalian epithelia. eLife, 2018. 7: p. e36739.

17. Maška, M., et al., The Cell Tracking Challenge: 10 years of objective benchmarking. Nature Methods, 2023. 20(7): p. 1010–1020.

18. Villars, A., et al., DeXtrusion: Automatic recognition of epithelial cell extrusion through machine learning <em>in vivo</em>. bioRxiv, 2023: p. 2023.02.16.528845.

19. Haubold, C., et al. A Generalized Successive Shortest Paths Solver for Tracking Dividing Targets. In Computer Vision – ECCV 2016. 2016. Cham: Springer International Publishing.

20. Guo, C., et al. On Calibration of Modern Neural Networks. 2017. 1706.04599 DOI: 10.48550/arXiv.1706.04599.

21. Çiçek, Ö., et al. 3D U-Net: Learning Dense Volumetric Segmentation from Sparse Annotation. 2016. 1606.06650 DOI: 10.48550/arXiv.1606.06650.

22. Platt, J.C., Probabilistic Outputs for Support Vector Machines and Comparisons to Regularized Likelihood Methods. 1999.

23. Mazzaferri, J., et al., Adaptive settings for the nearest-neighbor particle tracking algorithm. Bioinformatics, 2014. 31(8): p. 1279–1285.

24. Löffler, K., T. Scherr, and R. Mikut, A graph-based cell tracking algorithm with few manually tunable parameters and automated segmentation error correction. PLOS ONE, 2021. 16(9): p. e0249257.

25. Dietrich, F., Bayesian group belief. Social Choice and Welfare, 2010. 35(4): p. 595–626.

26. Dietrich, F. and C. List, 519Probabilistic Opinion Pooling, in The Oxford Handbook of Probability and Philosophy, A. Hájek and C. Hitchcock, Editors. 2016, Oxford University Press. p. 0.

27. Dey, T., et al., Survival analysis—time-to-event data and censoring. Nature Methods, 2022. 19(8): p. 906–908.

28. Aspert, T., D. Hentsch, and G. Charvin, DetecDiv, a generalist deep-learning platform for automated cell division tracking and survival analysis. Elife, 2022. 11.

29. Mohammadi, F., et al., A lineage tree-based hidden Markov model quantifies cellular heterogeneity and plasticity. Communications Biology, 2022. 5(1): p. 1258.

30. Murray, J.I., et al., Automated analysis of embryonic gene expression with cellular resolution in C. elegans. Nature Methods, 2008. 5(8): p. 703–709.

31. Matula, P., et al., Cell Tracking Accuracy Measurement Based on Comparison of Acyclic Oriented Graphs. PLOS ONE, 2015. 10(12): p. e0144959.

32. Ulicna, K., et al., Automated Deep Lineage Tree Analysis Using a Bayesian Single Cell Tracking Approach. Frontiers in Computer Science, 2021. 3.

33. Turetken, E., et al., Network Flow Integer Programming to Track Elliptical Cells in Time-Lapse Sequences. IEEE Transactions on Medical Imaging, 2016. PP: p. 1–1.

34. Moen, E., et al., Accurate cell tracking and lineage construction in live-cell imaging experiments with deep learning. 2019.

35. Hirsch, P., et al. Tracking by weakly-supervised learning and graph optimization for whole-embryo C. elegans lineages. 2022. 2208.11467 DOI: 10.48550/arXiv.2208.11467.

36. Wen, C., et al., 3DeeCellTracker, a deep learning-based pipeline for segmenting and tracking cells in 3D time lapse images. eLife, 2021. 10: p. e59187.

37. Stringer, C., et al., Cellpose: a generalist algorithm for cellular segmentation. Nature Methods, 2021. 18(1): p. 100–106.

38. Arbelle, A., et al., A probabilistic approach to joint cell tracking and segmentation in high-throughput microscopy videos. Medical Image Analysis, 2018. 47: p. 140–152.

39. Tian, C., C. Yang, and S. Spencer, EllipTrack: A Global-Local Cell-Tracking Pipeline for 2D Fluorescence Time-Lapse Microscopy. Cell Reports, 2020. 32: p. 107984.

40. Magnusson, K.E., et al., Global linking of cell tracks using the Viterbi algorithm. IEEE Trans Med Imaging, 2015. 34(4): p. 911–29.

41. Zheng, X., et al., Following cell type transitions in space and time by combining live-cell tracking and endpoint cell identity in intestinal organoids. bioRxiv, 2022: p. 2022.06.27.497728.

42. Krotenberg Garcia, A., et al., Active elimination of intestinal cells drives oncogenic growth in organoids. Cell Reports, 2021. 36(1): p. 109307.

43. Basak, O., et al., Induced Quiescence of Lgr5+ Stem Cells in Intestinal Organoids Enables Differentiation of Hormone-Producing Enteroendocrine Cells. Cell Stem Cell, 2017. 20(2): p. 177-190.e4.

44. McDole, K., et al., In Toto Imaging and Reconstruction of Post-Implantation Mouse Development at the Single-Cell Level. Cell, 2018. 175(3): p. 859-876.e33.

45. Verissimo, C.S., et al., Targeting mutant RAS in patient-derived colorectal cancer organoids by combinatorial drug screening. eLife, 2016. 5: p. e18489.

46. Barbáchano, A., et al., Organoids and Colorectal Cancer. Cancers (Basel), 2021. 13(11).

47. Lukonin, I., M. Zinner, and P. Liberali, Organoids in image-based phenotypic chemical screens. Experimental & Molecular Medicine, 2021. 53(10): p. 1495–1502.

48. Höfener, H., et al., Deep learning nuclei detection: A simple approach can deliver state-of-the-art results. Computerized Medical Imaging and Graphics, 2018. 70: p. 43–52.

49. Liu, R., et al. An Intriguing Failing of Convolutional Neural Networks and the CoordConv Solution. 2018. 1807.03247 DOI: 10.48550/arXiv.1807.03247.

50. Magnusson, K.E.G., et al., Global Linking of Cell Tracks Using the Viterbi Algorithm. IEEE Transactions on Medical Imaging, 2015. 34(4): p. 911–929.

51. Narayana, M. and D. Haverkamp. A Bayesian algorithm for tracking multiple moving objects in outdoor surveillance video. in 2007 IEEE Conference on Computer Vision and Pattern Recognition. 2007.

