## Supplementary figure for "Cell tracking with accurate error prediction"

### Supplementary figures

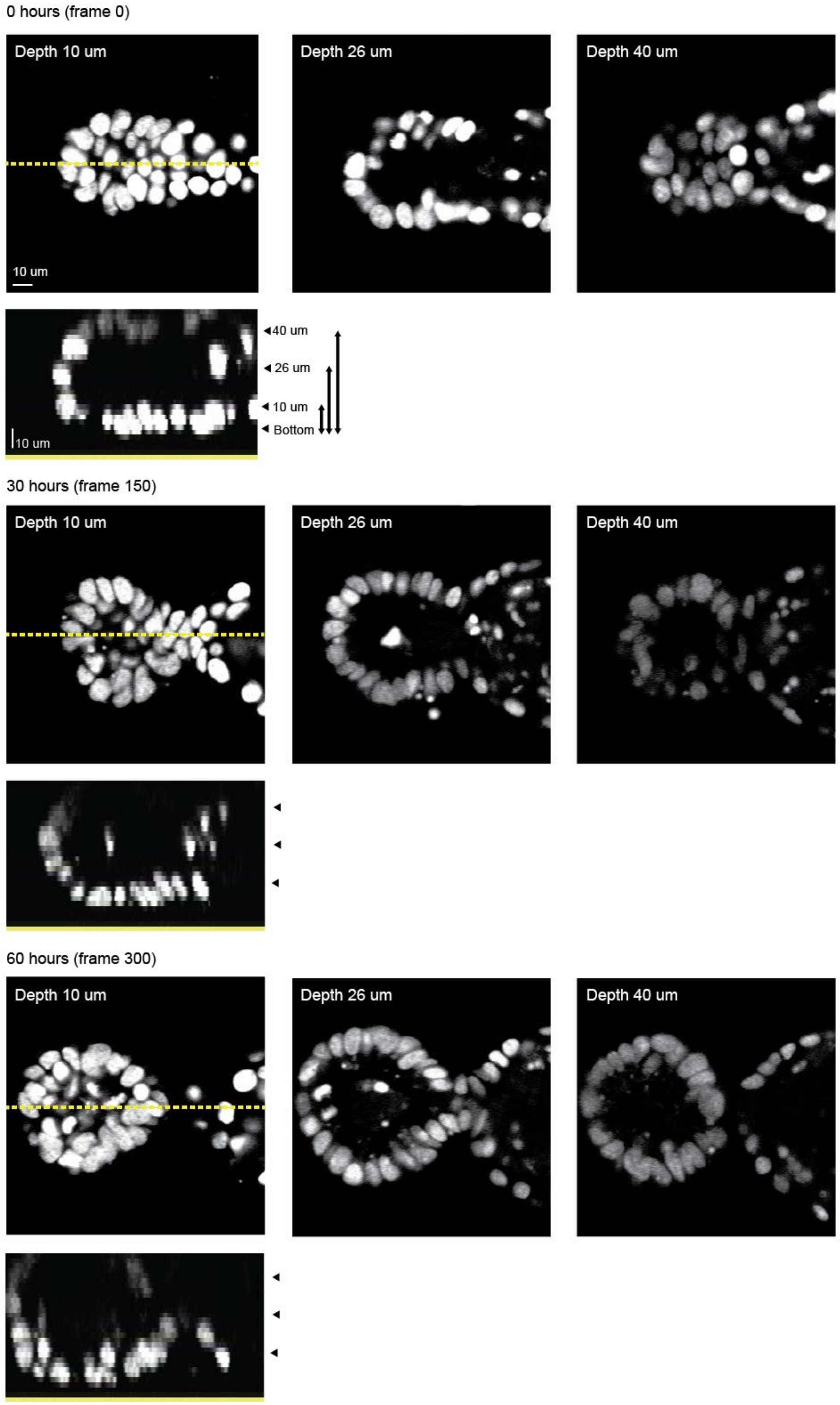


**Supplementary figure 1 - Overview of the data.** Images from a representative intestinal organoid used for training our neural networks. Z-planes are chosen relative to the position of the lowest nuclei. Nuclei are labeled using a H2B-mCherry construct. Nuclei are densely packed and can have strongly elongated shapes. Signal-to-noise levels are poor deep in the organoid and late in the experiment. The z-resolution (2 μm) is much lower than the xy-resolution (0.32 μm).

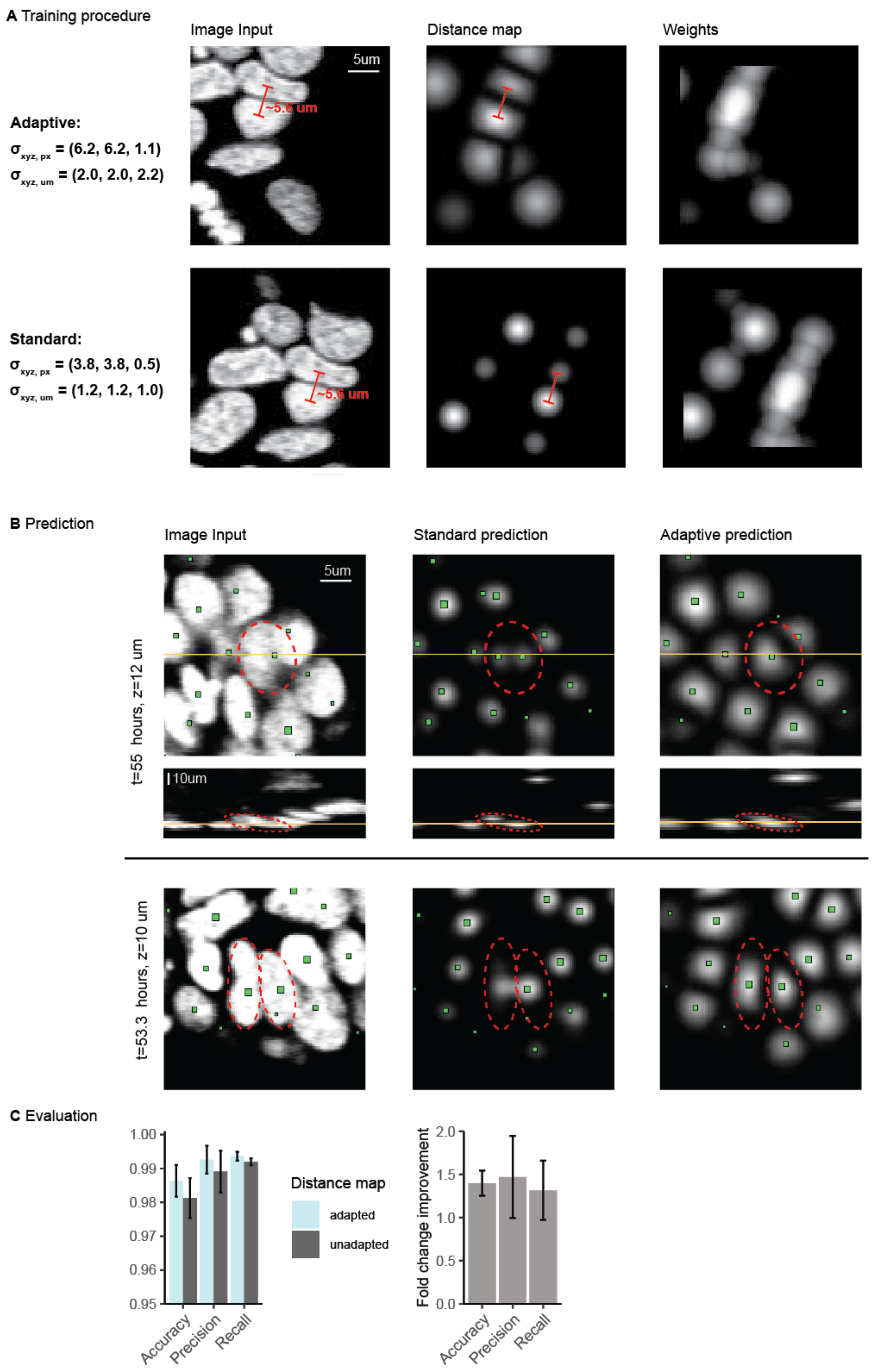


**Supplementary figure 2 – Adaptive distance mapping during cell center detection. A)** Distance maps for our adaptive method (top) versus the conventional way (bottom). The standard deviation of the Gaussian mapping of distances to intensity values are indicated by σ. They are shown in terms of pixels (px) and real distances (μm). For clarity we show training data generated from the same time point and chose overlapping crops to aid comparison. Shown are the input (first column), the desired outcome (second column) and the weight assigned to every pixel during the calculation of the loss (third column).

Our adaptive method allows the Gaussians around the cell centers to be much broader without overlapping, when in the conventional method Gaussians have to be narrower to not overlap. This is especially clear for non-spherical nuclei, where the centers can be very close together (see distance in red). The weights (third column) encode the contribution of a pixel towards the loss function. They are used to amplify the attention the neural network will pay to the nuclei containing regions relatively to the much larger background region. Gaussians for the weights can be chosen to be relatively broad to increase weights associated with the regions between nuclei, to ensure spots are well-separated. The edge regions, where nuclei are partially out of the field of view, receive zero weight. **B)** The performance of the adapted and non-adapted method on unseen test data. Examples are chosen to illustrate the improvements quantified in C). In the top row, the unadaptive method over-segments a large nucleus present in many z-slices, associating it with two Gaussian spots instead of single large one. In the bottom row, the Gaussian spots produced by the unadaptive method overlap, due to the closely packed and contorted nuclei, but is correctly separated in the adapted method. **C)** The false positive and negative rates for 5 different organoids using the adapted and the unadaptive distance mapping during training. Also shown is the fold chance improvement by using the adaptive distance mapping in false positive and negative rates for 5 different organoids. Error bars denote the standard deviation.

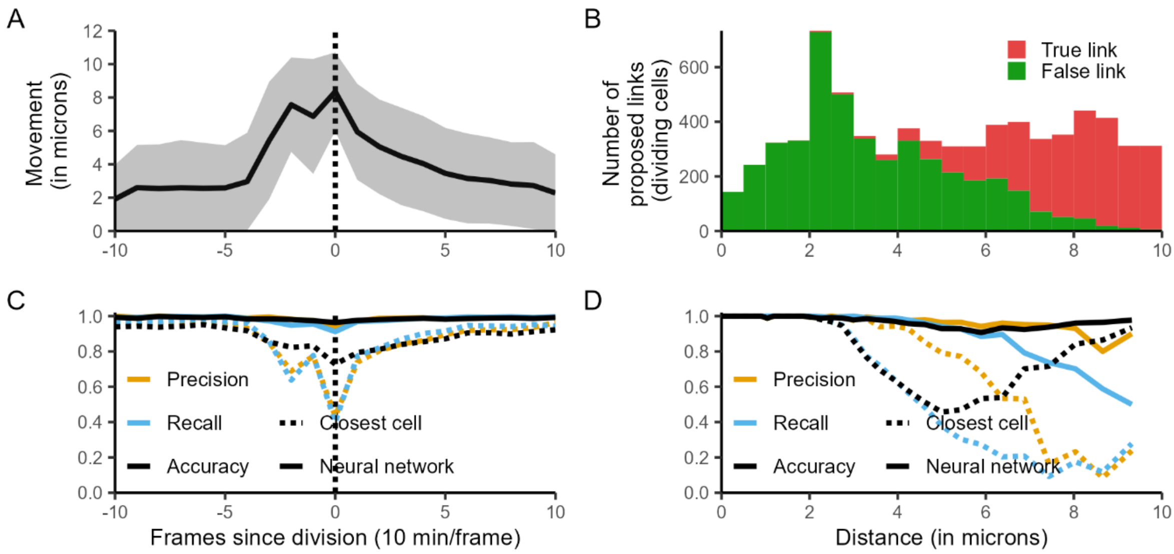


**Supplementary figure 3 - Link prediction for cells undergoing division. A)** The movement of nuclei per frame plotted against the frames since (or before) division (shaded region denotes standard deviation). Around the division, nuclei move substantially faster. **B)** Distribution of travel distance per frame for cells that are within 3 frames from the moment of division. All the proposed links in the graph are shown, whether true (green) or false (red). The distribution for true and false proposed links shows large overlap for dividing cells. **C)** Precision, recall and accuracy are shown as a function of frames since (or before) division. Naively picking the closest cell (dotted lines) breaks down as a mechanism for predicting links around division, when less than half of the true links is recalled (blue dotted line). In contrast, the neural network (solid lines) shows highly accurate prediction. **D)** Precision, recall and accuracy for the dividing subset of cells as a function of distance travelled by the nuclei. The neural network (solid lines) stays accurate (black line) even for fast movements, while linking the closest nuclei breaks down for dividing cell moving >3 μm in a single frame.

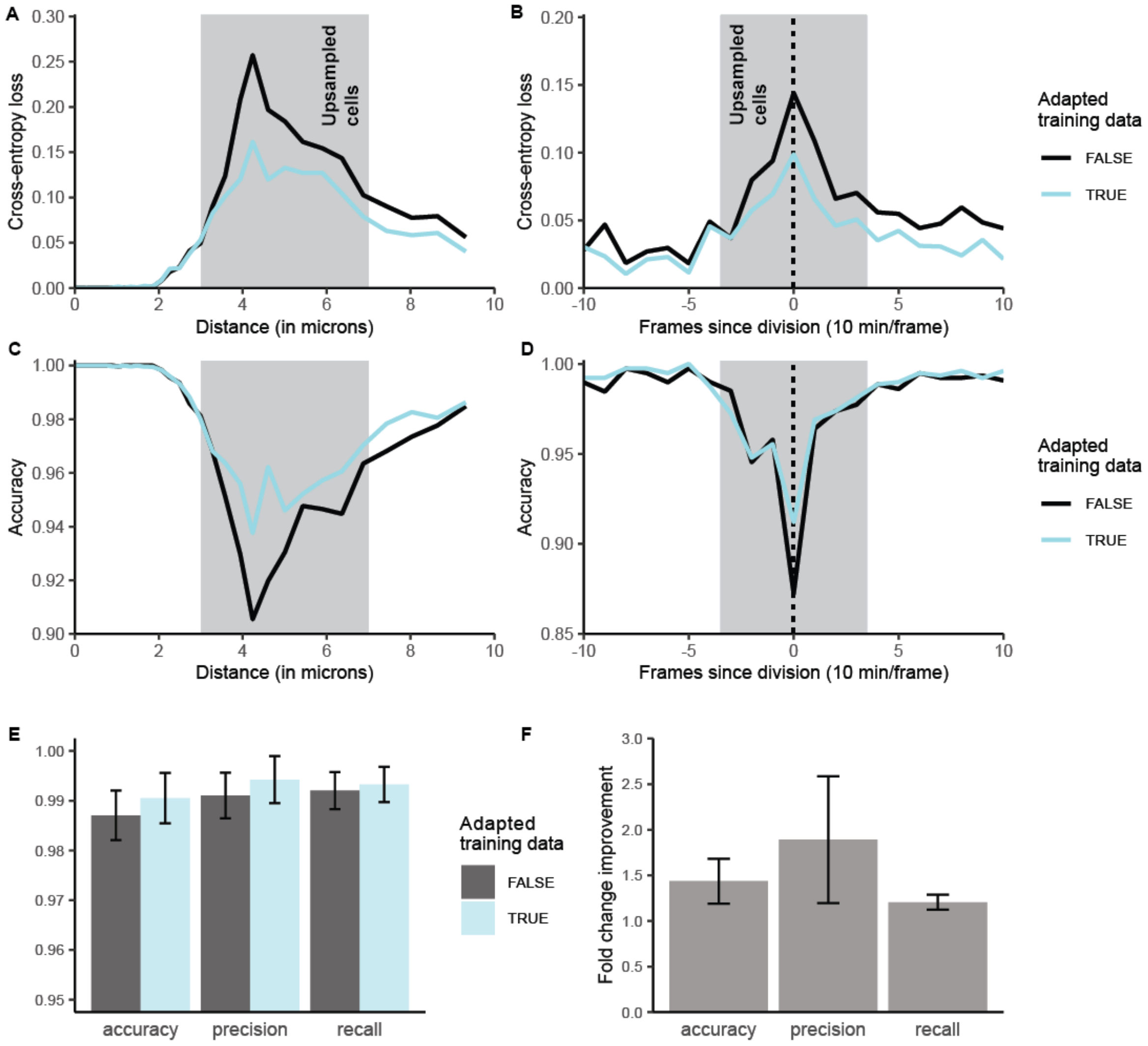


**Supplementary figure 4 - Impact of up-sampling the training dataset for link prediction. A, C)** The cross-entropy loss and accuracy for link prediction as a function of the distance between the two cell detections in the proposed link, for an up-sampled (blue) and regular training dataset (black). In the up-sampled training data challenging links with a distance of between 3 and 7 um (gray region) were increased by a factor of 5. Even small improvements in the cross-entropy loss and accuracy can lead to substantial improvements in tracking. The flow-solver that finds optimal tracks optimizes over all links, so improving the classification of a single link will help tracking on all nearby cells. Similarly, the error probabilities are calculated by integrating information from many links and thereby combine all the accuracy gains. **B, D)** The cross-entropy loss and accuracy for link prediction as a function of the time since (or before) division of the cell that forms the source node of the link. A five-fold up-sampling in the training data (blue) of cells close to division increases accuracy around the moment of division. **E)** Precision and recall before and after up-sampling the training data, for the complete dataset. Error bars shows the standard deviation of the 5 individual organoids that make up the test data**. F)** Fold change improvement in accuracy, precision and recall for every organoid in the test data upon up-sampling the training data. Error bars denote the standard deviation between different organoids.

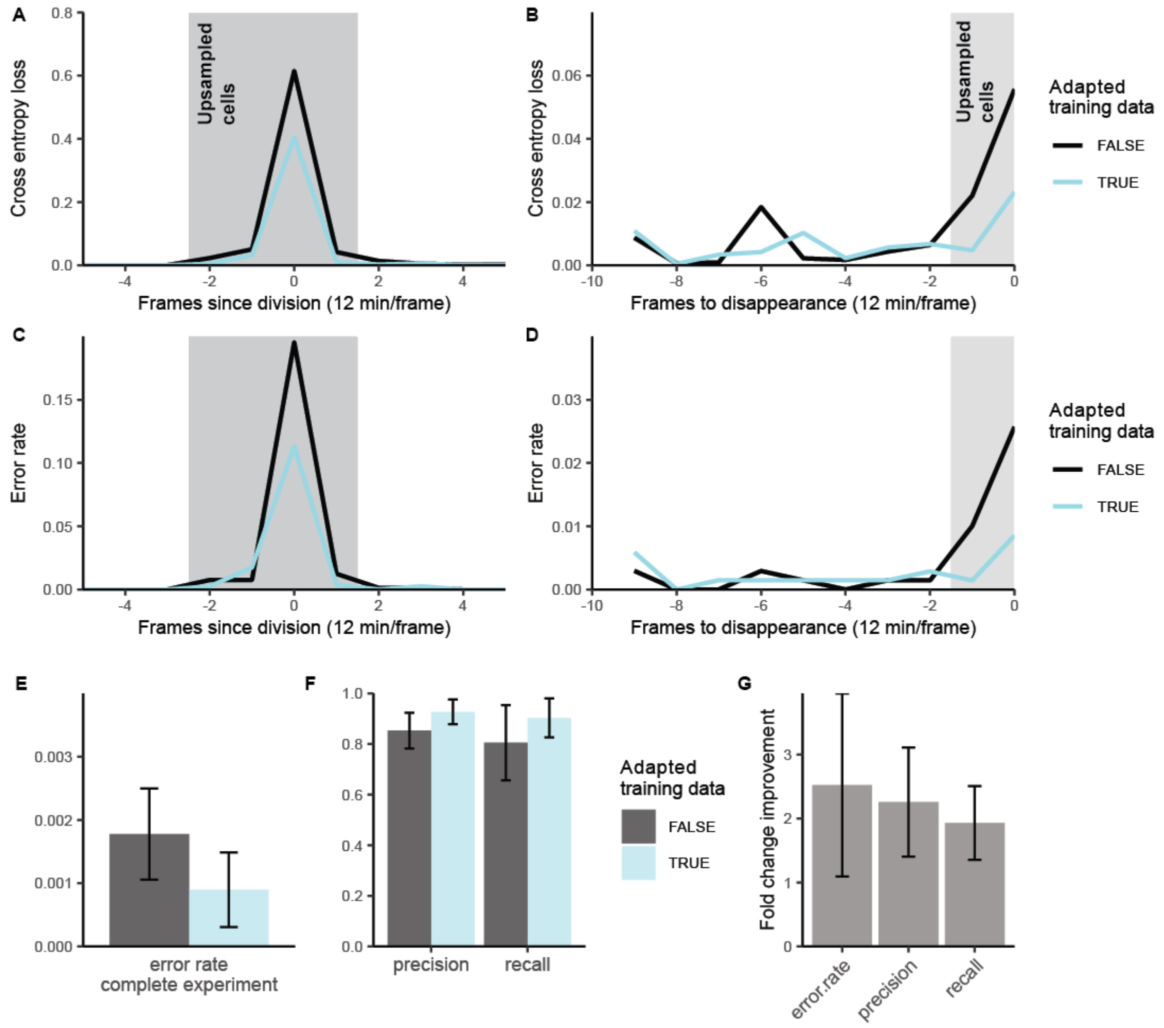


**Supplementary figure 5 - Impact of up-sampling the training dataset for division prediction. A, C)** The cross-entropy loss and error rate for cell division prediction around the moment of division for an up-sampled (blue) and regular training dataset (black). The up-sampled training data increased cells around division (gray region) 10-fold. **B, D)** The cross-entropy loss and error rate for cell division prediction as function of time prior to the end of a cell track. The majority of cell tracks end due to cell death. As dying cells resemble dividing cells, the error rate without up-sampling increases to almost 3% (black line). For up-sampled training data (blue line), the error rate only increases for the final time point before death and only to 1% **C)** Error rates for all cell detections in the test data with (blue) and without up-sampling the training data (gray). Error bars shows the standard deviation of the 5 individual organoids that make up the test data. **E)** Precision and recall measures with and without up-sampling the training data. **F)** Fold change improvement in error rate, precision and recall for every organoid in the test data upon up-sampling the training data. Error bars denote the standard deviation between different organoids. Both the amount of false positives (precision) and false negatives (recall) decreases around two-fold by up-sampling the training data.

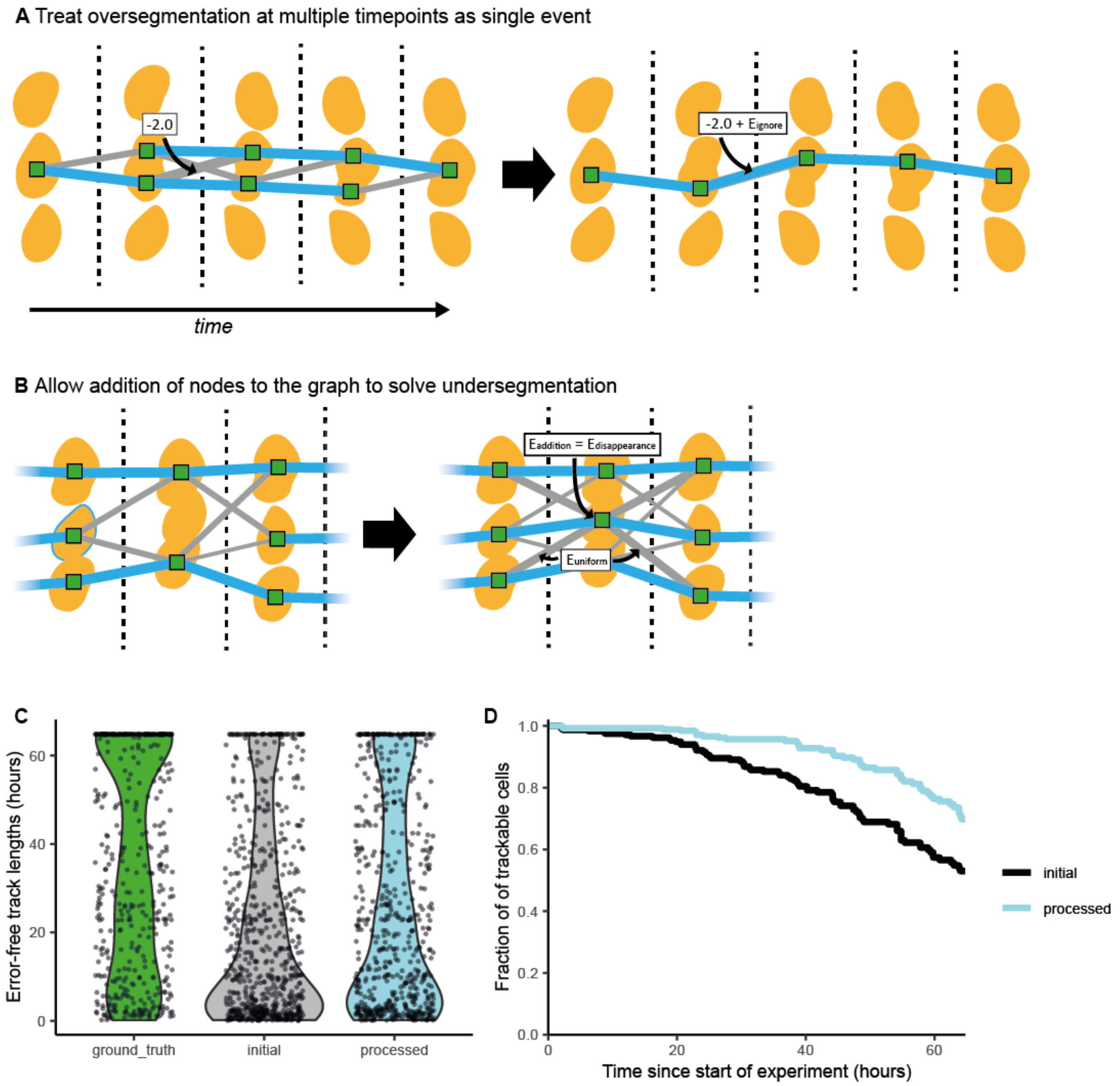


**Supplementary figure 6 - Post-processing of the maximum likelihood tracking solution. A)** Illustration of how post-processing deals with over-segmentations. The blue links represent the maximum-likelihood solution, while the grey links are excluded from the solution. Over-segmentations are not independent events, but are treated as such by the min-cost flow solver, which can lead to fragmentation of tracks (left panel). During post-processing we check if track fragments can be joined by adding a single link and removing the cell detections associated with the over-segmentation. Removing multiple of these nodes is associated with a single penalty ($E_{ignore}$, the relative log-likelihood of over-segmentation) which is added to the new link so that our probabilistic description remains correct. **B)** Illustration of how post-processing deals with under-segmentations. The min-cost flow solver cannot add nodes to the graph, which can lead to fragmentation of tracks (left panel). During post-processing we check if track fragments can be joined by adding a node. Adding a node is associated with a penalty ($E_{addition}$, the relative log-likelihood of missing a cell detection) and the node is connected to nearby nodes in the graph with uniform probability. **C)** Error-free track length distributions in a representative tracked organoid. The ground truth (green) represents fully manually corrected data, where tracks are only cut short by cell death, the end of the experiment, or leaving the field-of-view. The initial maximum-likelihood solution (gray) has many more short tracks, which is partly solved by post-processing (blue). **D)** The fraction of cells present at a certain time that can be tracked without error from the start of the experiment. Only cells that are trackable for the full experiment in the ground truth are considered. By post-processing the data we can increase the fraction of cells that are trackable for the complete experiment (> 60 hours) from around 50% to around 75%.

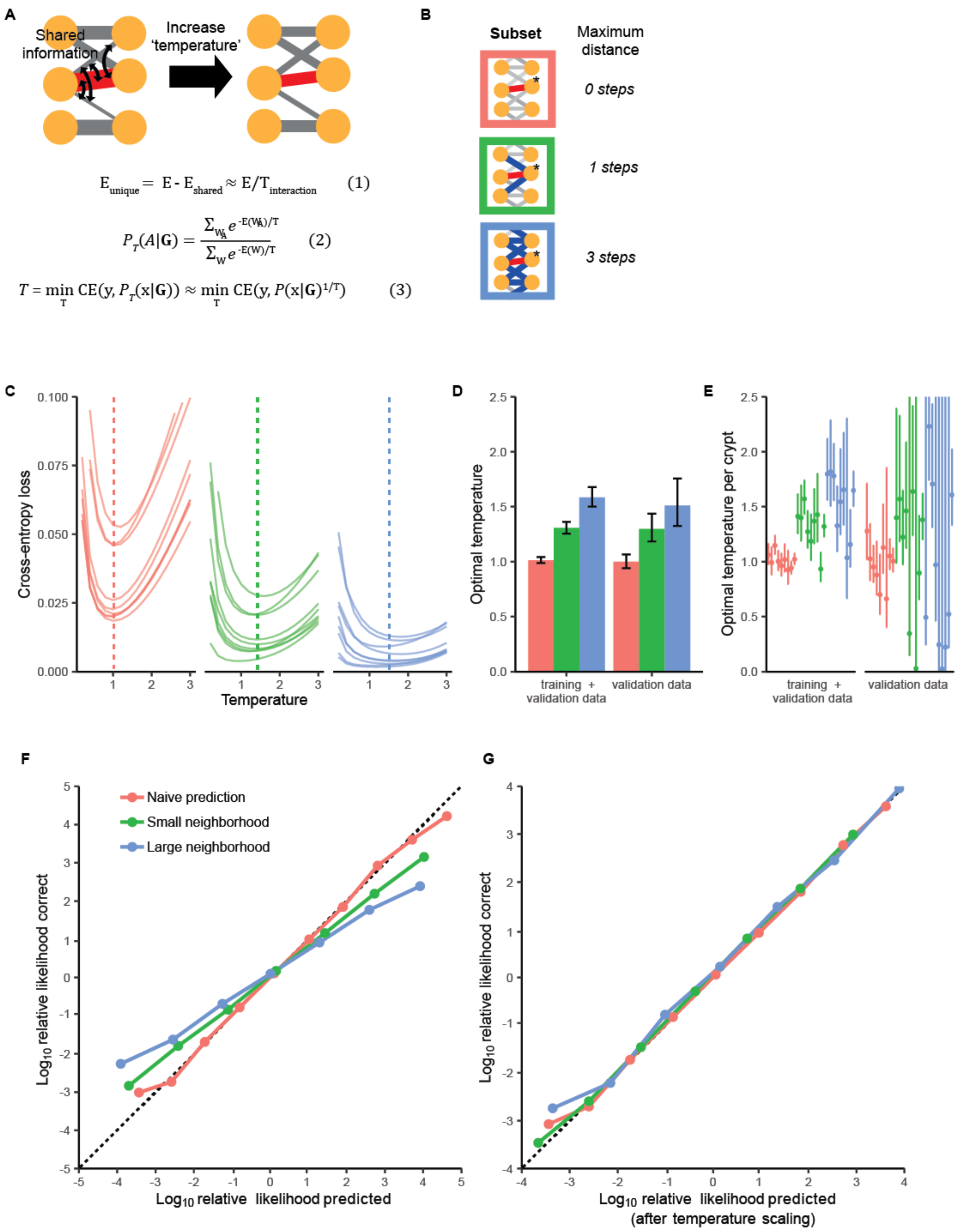


**Supplementary figure 7 - Temperature scaling during the marginalization procedure. A)** Visual illustration of the temperature scaling procedure. The prediction on a single link (encoded in its weight) is not independent from predictions on nearby links. To account for this shared information we divide the link energy by a 'calibration temperature' (Eq. 1). The marginalized link probability, P(A|B), is then based on scaled energies (Eq. 2). The calibration temperature is found by minimizing the cross-entropy loss (CE) between the marginalized predictions and the ground truth (y), with respect to this temperature (Eq. 3). This scaling temperature has to be calibrated only once after training a set of division and link neural networks. The calibration can be done on the same data that was used for neural network training. For data far outside the training distribution new calibration can be performed on a small manually corrected set of links. **B)** Illustration of the difference subsets used in the subsequent plots. Red symbolizes the naïve approach, where the subset simply is the link of interest. In this case no marginalization is done. The green subset only considers link at a distance of one step from the target node of the link of interest. The blue subset goes up to a distance of three steps. This is the largest set that is computationally feasible (~ 1 hour of computation time for 300 frames). **C)** Cross entropy loss between predictions and ground truth (based on all 9 organoids in the training dataset) for different neighborhoods and temperatures. Minimum loss (dotted line) is achieved at higher temperatures when the subset gets larger. All lines represent individual organoids. **D)** Optimal calibration temperature for every neighborhood. The error bars represent the 95% confidence interval (this is a lower bound as not all observed links are truly independent). Limiting ourselves to calibration on the validation dataset that was left during training gives the same results. **E)** Optimal temperatures and confidence intervals (again lower bounds) per organoid. The estimates show great overlap between organoids, validating that we can use a single calibration temperature for all. **F)** Predicted versus actual log-likelihoods after marginalization on different subsets. When marginalizing on larger subsets predictions become overconfident. If predictions suggest that links are not correct, more than expected fraction is actually correct and the other way around. **G)** After scaling the energies with the proper temperature per subset, we get well-calibrated link predictions.

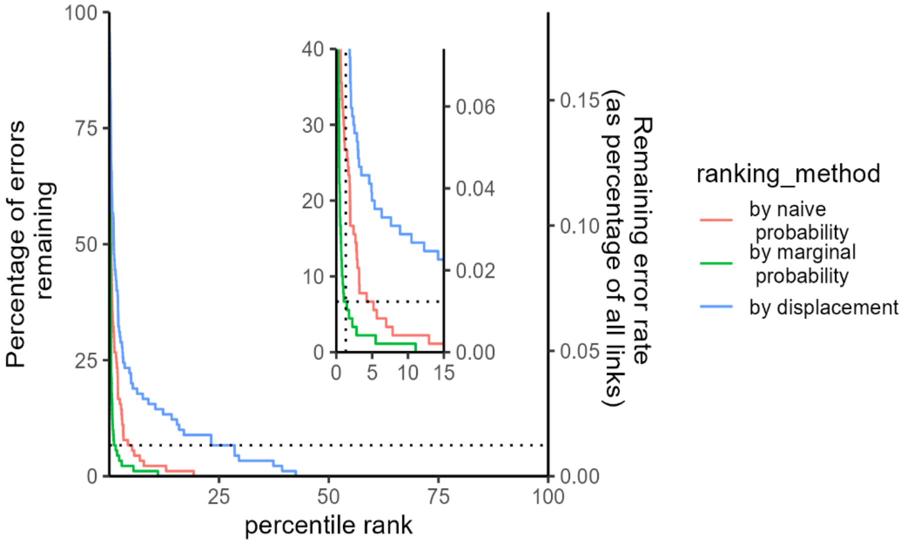


**Supplementary figure 8 – Error rates versus heuristic.** The comparative performance of marginalized error rates against naïve error rates and a typical heuristic for linking errors, distance travelled, is shown. If all links with an error rate of above 1% are reviewed, >90% of the errors in the data are corrected (horizontal dotted line). This leads to a remaining linking error rate of at most ~ 0.01% per cell per time point. This is an upper bound as review might also correct nearby errors that above the confidence threshold. To achieve this, the user has to check only the ~1% unlikeliest links in the data (vertical dotted line in inset). Using the naive neural network predictions, this would require checking ~5% of the data (red curve in inset). Without probabilistic information, a user would have to manually check the ~25% biggest displacements in the dataset to reach comparable accuracy (blue curve).

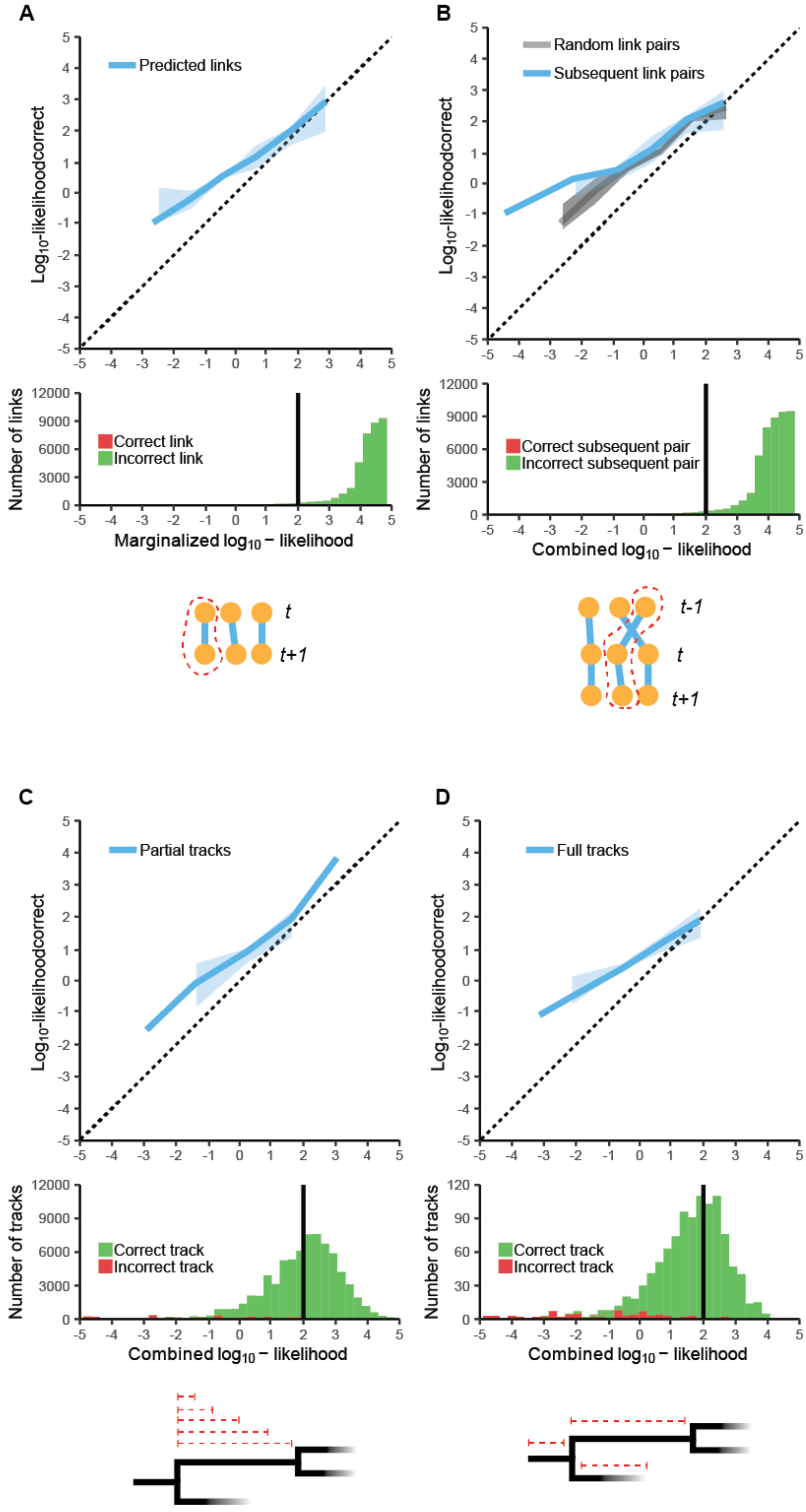


**Supplementary figure 9 – Track error rates. A)** The predicted likelihood (after marginalization) of links in the tracking solution against the measured likelihood of being correct based on the manual data (top panel). The line does not overlap the dotted line, because not all graph information available to the flow-solver can be integrated in the error prediction. Key is that they remain above the dotted line so that the error predictions are conservative. The histogram (bottom panel) shows the distribution of likelihoods of correct (green) and incorrect links (red). Because the linking error rate is so low, incorrect instances cannot be seen in the histograms A) and B). The black vertical line indicates the threshold of 99% chance of being correct. **B)** The likelihood of a pair of links both being correct can be calculated by combining their constituent probabilities by simple multiplication. It does not matter if the links are subsequent (blue line) or unconnected (gray line). It is thus not so that a link being true is informative of the truth of the subsequent link, beyond its predicted error probability. **C)** The predicted error rates for tracks of arbitrary length. The probability that a cell can be correctly traced back to its last division (red dotted lines) is predicted for every cell at every timepoint. The probability of the track being correct is calculated by multiplying all the constituent probabilities. The tracks are compared to the ground truth and deemed correct if they recapitulate it exactly, yielding a similar calibration curve to A). **D)** The same as in C) but now only with tracks that span the full cell cycle, again producing a similar calibration curve as in A). Error detection works efficiently for full tracks spanning the complete cell cycle, only one incorrect track is above the 99% certainty cut-off.

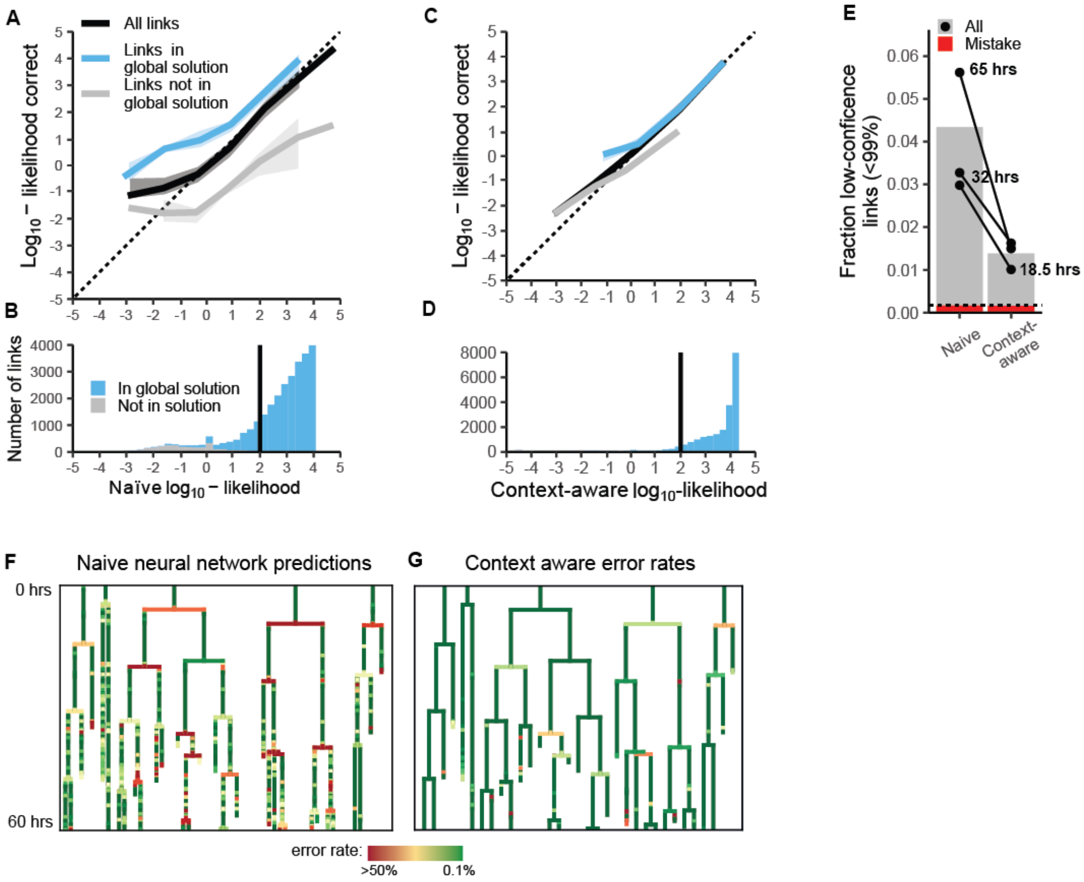


**Supplementary figure 10 - Marginalization during full procedure. A)** The predicted relative $\log_{10}$-likehoods from the link neural network (black line) is well-calibrated when compared to the manually corrected ground truth (dotted line denotes perfect calibration). The overestimation at low likelihoods is due to the filtering of unlikely links before tracking and marginalization (see **Methods**). This filtering preferentially removes low-probability incorrect links, making the remaining ones more likely to be true. The fact that links in the global solution (blue line) are much more likely than expected and links excluded from the tracking solution (gray line) are less likely than expected, suggests that contextual information could improve the error rate prediction. **B)** Many links in the tracking solution are less than 99% (black line) certain based on the naïve predictions. **C)** Marginalization integrates context and largely removes the discrepancy between links in and out of the tracking solution. **D)** Only few links in the tracking solution are less than 99% certain after marginalization. **E)** The fraction of uncertain links (<99% certainty) as fraction of total. Marginalization decreases the amount of uncertain links around four-fold. The longest experiments, which have poorer signal to noise, benefitted most from the marginalization (the black lines denote individual experiments). The red fraction indicates actual errors in the low-confident fraction. The dotted line the fraction of errors across all links including ones that have a high probability of being true, showing that no significant amount of errors is missed. **F)** Five randomly selected lineage trees colored by their naïve predicted error rate (yellow to red links are uncertain. **G)** The same trees after marginalization.

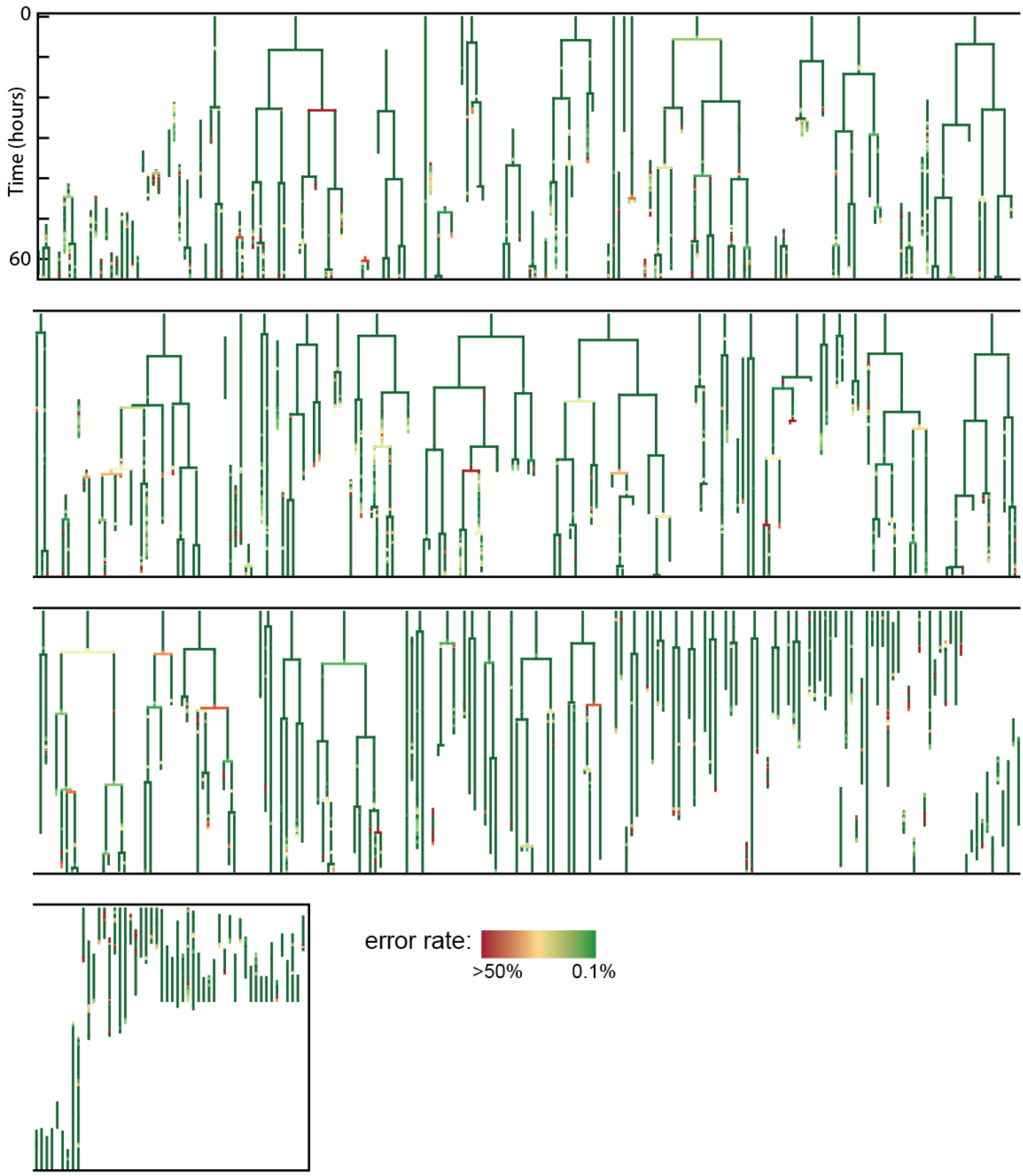


**Supplementary figure 11 – All trees with error rates before correction for typical organoid.** All cell tracks over 5 hours long for a single organoid with one crypt plus villus region imaged. Data is shown before any correction and with the context-aware error rates. Note that many cells can already be traced back to the start going only through high-confidence (green) links.

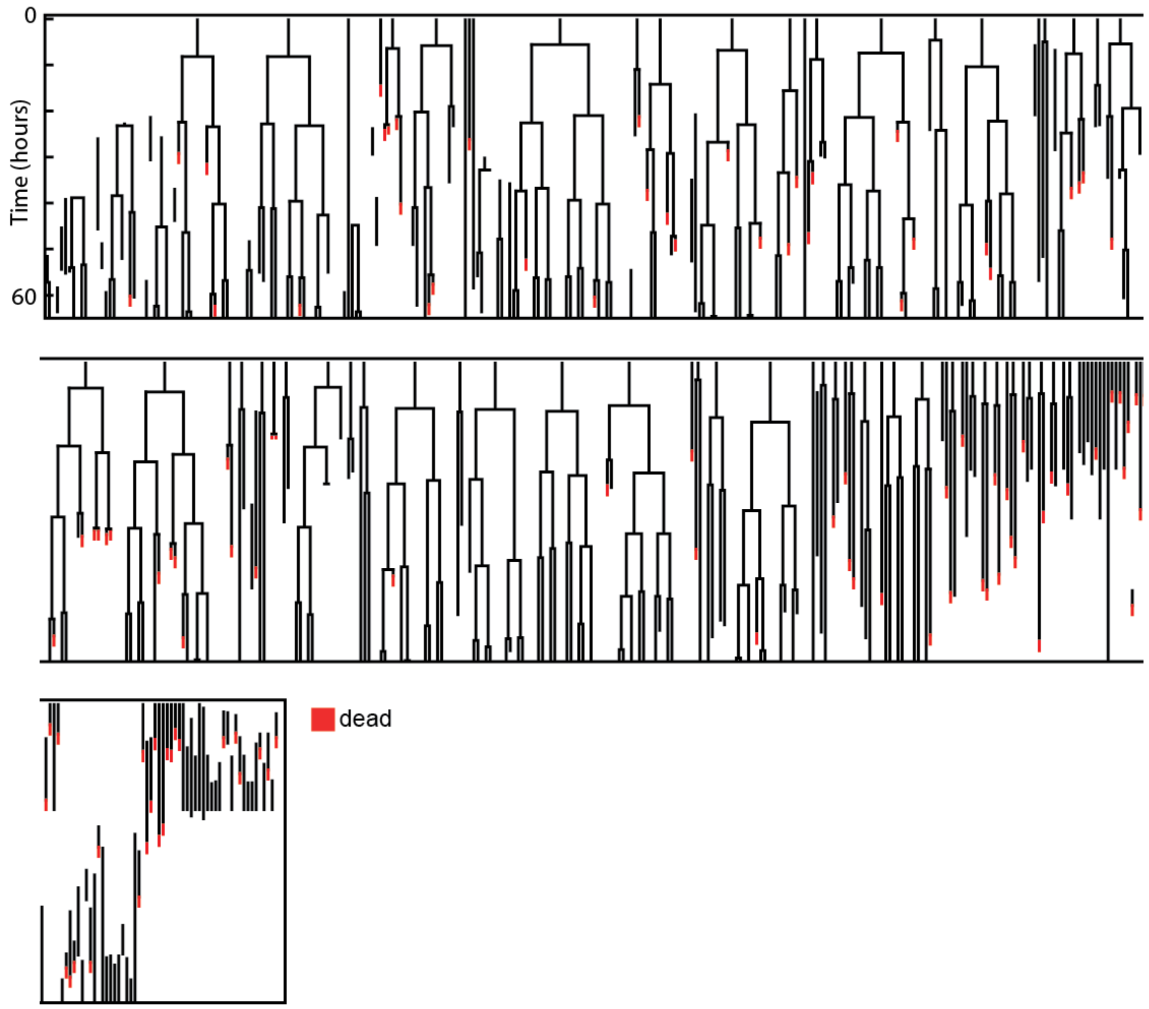


**Supplementary figure 12 – All trees with error rates after correction for typical organoid.** All cell tracks over 5 hours long for a single organoid with one crypt plus villus region imaged. All track that end and are not marked 'dead' leave the field of view.

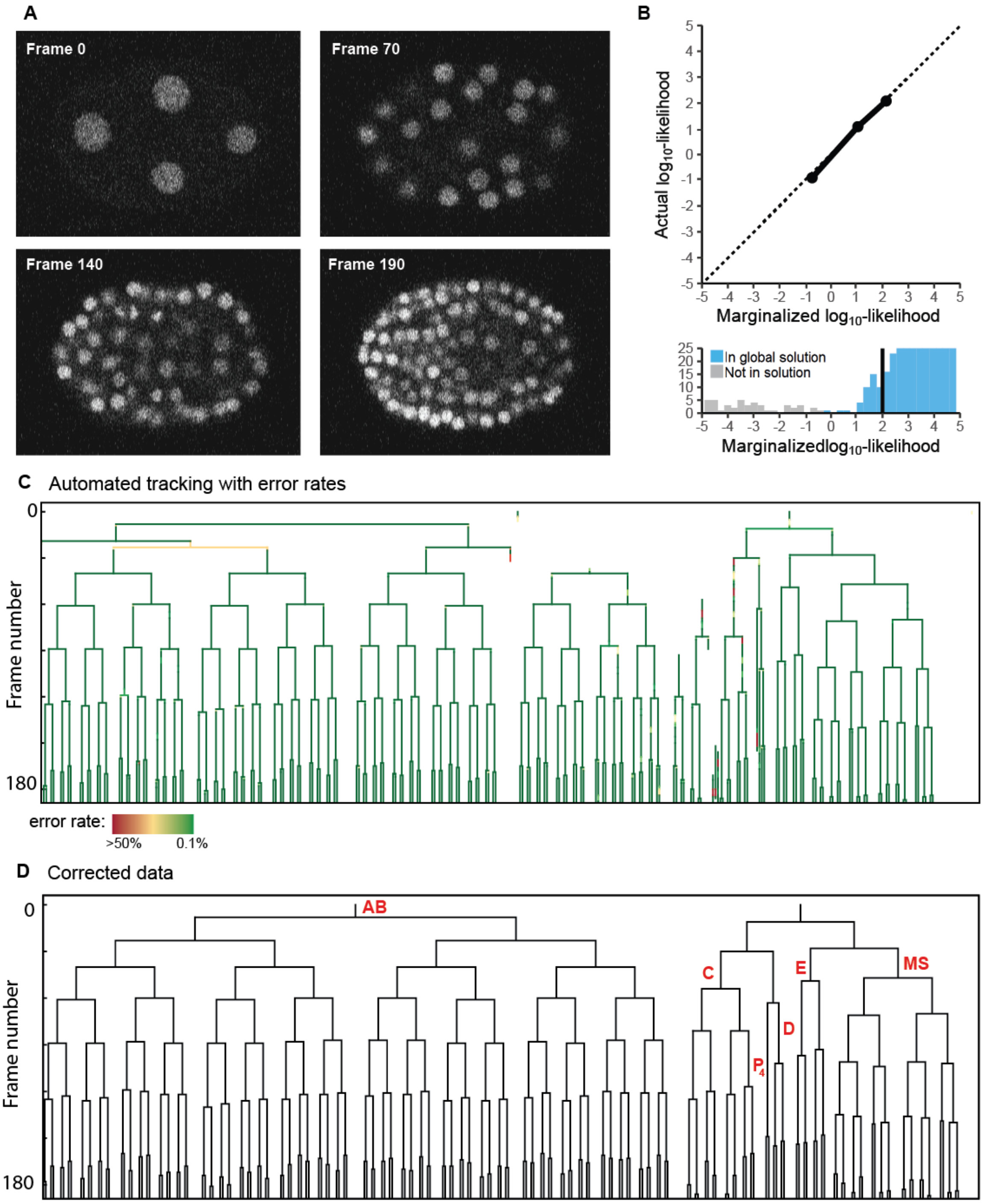


**E** Tracking statistics

| | count | rate |
|---|---|---|
| *Total cell detections* | 11045 | - |
| *Wrong links* | 3 | .03% |
| *Oversegmentation* | 3 | .03% |
| *Undersegmentation* | 32 | .29% |

**Supplementary figure 13**: **A)** Z-slices of 3D confocal data for *C. elegans* embryogenesis (training data). **B)** Using retrained neural networks, we obtain well-calibrated link-likelihood predictions (top panel). Almost all links in the tracking solution that minimize the global energy (blue) have very high marginalized likelihoods (bottom panel). In this data set, only few links in the graph are not in the tracking solution (grey) as the nuclei are less closely packed then in intestinal organoid data. **C)**

Lineage trees with associated error rates show that only few potential errors (41 links with an error probability above 1%) are present after automated tracking. **D)** Lineage trees after manual correction of potential errors. The lineages map exactly on the known *C. elegans* AB, E, MS, C, D and $P_4$ sub-lineages, strongly suggesting that no errors remain. **E)** Tracking statistics show a very low error rate and that almost no error in linking. Most errors arise from undetected nuclei (under-segmentation), deep in the imaging volume.

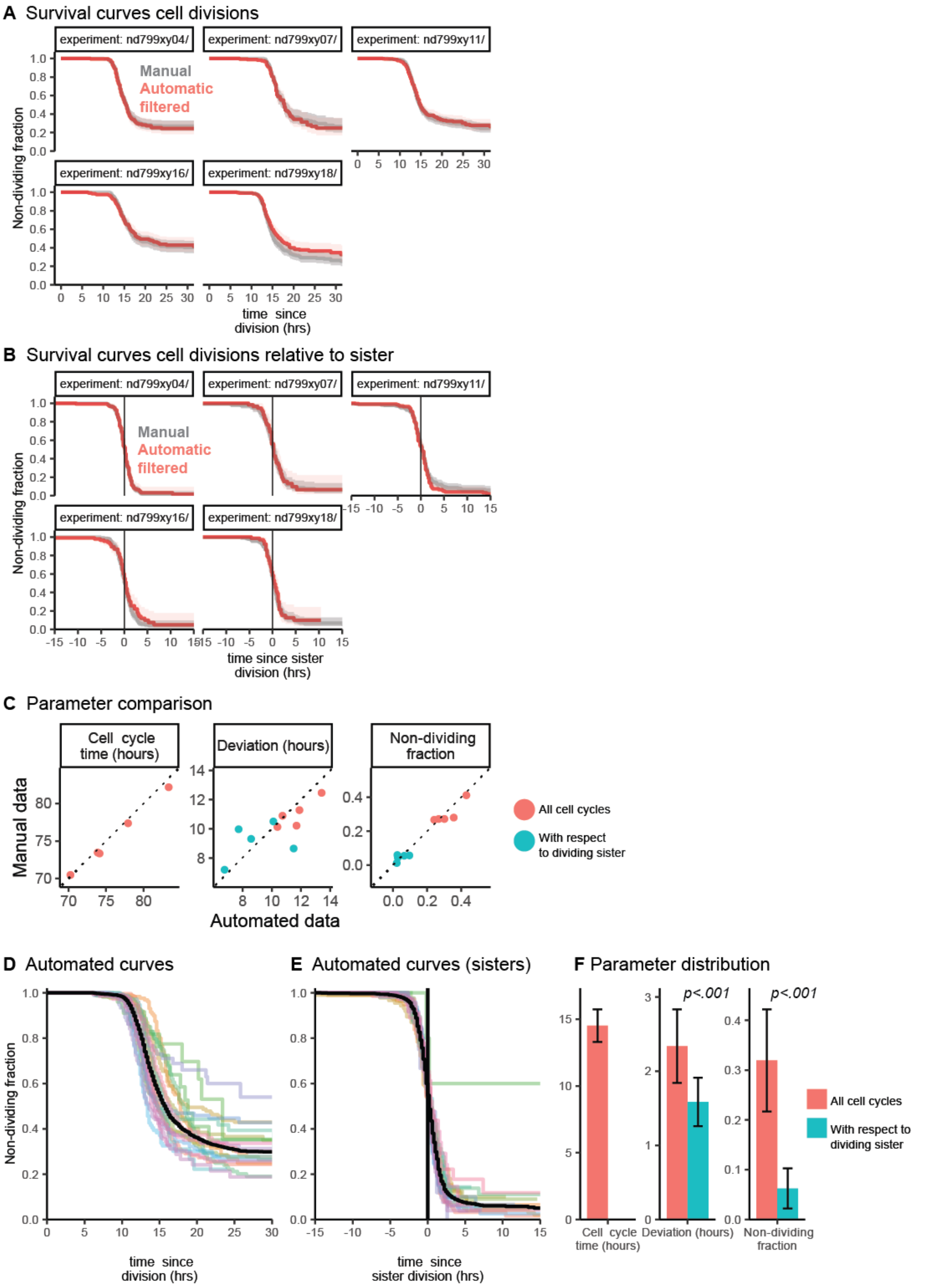


**Supplementary figure 14 – Survival curves for different organoids. A)** Kaplan-Meier curves describing the fraction of non-divided cells as a function of time since division, for 5 organoids where manual reference data was available for the complete crypt (grey curve). The shaded region denotes the 95% confidence interval. **B)** Kaplan-Meier curves describing the fraction of non-divided cells as a function of the time since the division of the sister cell. **C)** Comparison of manually annotated data to automated analysis for the estimation of three key parameters of lineage dynamics. Color indicates whether the variation in cell cycle duration or the probability that the cell will divide again was calculated for all cell cycles (red) or for cell cycles relative to the moment of division of the sister cell (blue). Cell cycle times show almost perfect correlation between manual and automated data. The deviation in the cell cycle time is more sensitive to outliers and consequently shows poorer correlation. The non-dividing fraction again shows strong correlation. **D)** Overlay of all Kaplan-Meier curves of automatically tracked organoids, n=20, which describes the fraction of non-divided cells as a function of time since division. Black line represents the combined data. **E)** Same but as a function of time since the sister division. **F)** Statistical analysis of organoid lineage parameters. Error bars indicate standard deviations around the mean. Cell cycle times only show limited variation across organoids and experiments, with standard deviation <10% of the mean cell cycle time (left panel). The variation around the cell cycle mean for cell cycle times is significantly larger than the variation between the cell cycle times of sisters (middle panel), suggesting strong correlation between sister pairs. This is further supported by the results that given that the sister divides, the non-dividing fraction becomes close to zero instead of around 30% (right panel).

### Supplementary text

**Individual-wise bayesianeity**

Dietrich and List motivate multiplicative opinion pooling by showing it conforms to a strong intuition: that it should not matter if information is presented after the pooling procedure or beforehand to an individual predictor. They term this axiom 'individual-wise bayesianeity' (Dietrich, 2010; Dietrich & List, 2016). Following their notation we introduce a likelihood function $L(a)$, which encodes the extra information about an element $a$, such that updating our prediction after pooling would give:

$$P^L(a) = \frac{P(a)L(a)}{1 - P(a) + P(a)L(a)} \tag{1}$$

Assume now that this information is known beforehand to the neural network (an individual predictor) assigned to predict event $a$. The consensus probability would then be given by:

$$P^L(a) = \frac{\sum_{\omega \ni a} L_{0,\omega}\, L(a)\, \prod_{i \in \omega} \frac{L_i}{L_{i,prior}}}{\sum_{\omega \setminus\ni a} L_{0,\omega} \prod_{i \in \omega} \frac{L_i}{L_{i,prior}} + \sum_{\omega \ni a} L_{0,\omega}\, L(a) \prod_{i \in \omega} \frac{L_i}{L_{i,prior}}} \tag{2}$$

$$P^L(a) = \frac{L(a) \sum_{\omega \ni a} L_{0,\omega} \prod_{i \in \omega} \frac{L_i}{L_{i,prior}}}{\sum_{\omega} L_{0,\omega} \prod_{i \in \omega} \frac{L_i}{L_{i,prior}} + (L(a) - 1) \sum_{\omega \ni a} L_{0,\omega} \prod_{i \in \omega} \frac{L_i}{L_{i,prior}}} \tag{3}$$

If we then numerator and denominator by the partition function ($\sum_{\omega} L_{0,\omega} \prod_{i \in \omega} \frac{L_i}{L_{i,prior}}$), we get:

$$P^L(a) = \frac{L(a)P(a)}{1 + (L(a) - 1)P(a)} \tag{4}$$

$$P^L(a) = \frac{P(a)L(a)}{1 - P(a) + P(a)L(a)} \tag{5}$$

Which is the same as equation (1) indicating that it does not matter at what stage information is introduced in our procedure and we conserve the 'individual-wise bayesianeity' axiom. It also provides us with a convenient way to incorporate new information about an element without redoing marginalization.

More generally we can see that integrating extra information about an element $i$ during opinion pooling regarding any element $a$, is equivalent to updating the probability the following way after opinion pooling:

$$P^L(a) = \frac{P(a,i)L(i) + P(a,\neg i)}{1 - P(i) + P(i)L(i)} \tag{6}$$

This is in practice not a useful way of incorporating post hoc information as $P(a,i)$ and $P(a,\neg i)$ have to be computed using our marginalization approach as well.

In our derivations above we assume that the extra information shares zero overlap with the previously available information. If this would not hold we should discount $L(a)$ by dividing it by an appropriate temperature as proposed in the main text and method section, but this would not affect our argument.

**Bayesian belief matrix**

Another way of integrating multiple probabilities is using a Bayesian belief matrix approach (Ulicna, Vallardi, Charras, & Lowe, 2021). Here information is integrated sequentially using Bayesian update rules. Per integrated prediction the process has two steps. First we update the belief about the element the prediction is made on:

$$P_{n+1}(a) = \frac{P_n(a)L(a)}{1 - P_n(a) + P_n(a)L(a)} \tag{7}$$

We then normalize all other probabilities to satisfy the constraints of the tracking problem. This method is limited in which type of constraints it can take into account, because all constraints have to be implemented in a single normalization step. It can therefore only deal with mutually exclusive events, such a links connecting towards same node under the condition that cell merging is prohibited. For every event $i$ mutually exclusive with element $a$, the updated probabilities become:

$$P_{n+1}(i) = P_{n+1}(i)\frac{1 - P_{n+1}(a)}{1 - P_n(a)} \tag{8}$$

If we then substitute $P_{n+1}(a)$:

$$P_{n+1}(i) = P_{n+1}(i)\frac{1 - \dfrac{P_n(a)L(a)}{1 - P_n(a) + P_n(a)L(a)}}{1 - P_n(a)} \tag{9}$$

$$P_{n+1}(i) = P_{n+1}(i)\frac{\dfrac{1 - P_n(a)}{1 - P_n(a) + P_n(a)L(a)}}{1 - P_n(a)} \tag{10}$$

$$P_{n+1}(i) = \frac{P_{n+1}(i)}{1 - P_n(a) + P_n(a)L(a)} \tag{11}$$

This is identical to equation (6) for mutually exclusive events $i$ and $a$, in which case $P(a, i)$ is zero and $P(a, \neg i)$ is the same as $P(a)$. We can thus see that the 'Bayesian belief matrix' update rules are simply the same as the rules for integrating information post-hoc in our multiplicative pooling framework. 'Individual-wise bayesianeity' then guarantees, that the stepwise updating of via Bayesian update rules is mathematically equivalent to our marginalization approach in the case of mutually exclusive events. The starting probabilities $P_0(i)$ function in this case as the priors.

Our approach can even be called a generalization in the sense that the Bayesian updates can only deal with elements that are mutually exclusive, otherwise the normalization step would not work. While our method can also deal with more advanced constraints, such as that two links can exit a node but only if the cell is dividing.

Note that our description here differs slightly from the one given by Ulicna et al.. In their method there are no neural networks estimating probabilities of events from the observed data. Instead they work with models that give the probability of observing data given the event we want to predict. Their version of equation (7) would thus take the form of:

$$P_{n+1}(a) = \frac{P_n(a)\dfrac{P(\theta_a|a)}{P(\theta_a|\neg a)}}{\big(1 - P_n(a)\big) + P_n(a)\dfrac{P(\theta_a|a)}{P(\theta_a|\neg a)}} \tag{8}$$

In which the likelihood information $L(a)$ is thus of the form:

$$L(a) = \frac{P(\theta_a|a)}{P(\theta_a|\neg a)} \tag{9}$$

The probability of observing the data when an event (like a link) is not true $P(\theta_a|\neg a)$ in these type of methods and are not used directly. Instead this probability can be assumed to be equal for all events in the system. The fact that all events are mutually exclusive then allows $P(\theta_a|\neg a)$ to be simply multiplied out:

$$P_{n+1}(a) = \frac{P_n(a)P(\theta_a|a)}{\big(1 - P_n(a)\big) + P_n(a)P(\theta_a|a)} \tag{10}$$

This assumption is reasonable for similar events (different links), but clearly not when considering for instance the chance of a cell disappearing $P(d)$. This problem is ameliorated by Ulicna et al. by capturing it in a single hyper parameter, $P^*(\theta|d)$, by which to divide all link likelihoods:

$$P^*(\theta|d) = \frac{P^*(\theta|d)P(\theta_a|\neg a)}{P^*(\theta|\neg d)} \tag{11}$$

This factor integrates the chances of cell disappearances in the predictions, without needing to explicitly update them:

$$P_{n+1}(a) = \frac{P_n(a)P(\theta_a|a)}{\left(1 - P_n(a)\right)P^*(\theta|d) + P(a)P(\theta_a|a)} \tag{12}$$

The fact that $P(\theta_a|\neg a)$ is not used directly reflects the fact that it is difficult to define models for these type of probabilities. This highlights an advantage of neural networks that instead aim to estimate the probability of an event given the data $P(a|\theta_a)$. The likelihood information then simply becomes:

$$L(a) = \frac{P(\theta_a|a)}{P(\theta_a|\neg a)} = \frac{P(a|\theta_a)P(\neg a)}{P(\neg a|\theta_a)P(a)} = L_\theta(a)/L_{prior}(a)$$

In which an uniform prior could be chosen like in Ulicna et al. or it can be chosen to reflect possible shared information as described in the methods section. The final outcome of the 'Bayesian belief matrix' approach can then simply be calculated as (with $L(d)$ giving the likelihood of a cell disappearing):

$$P_{final}(a) = \frac{L(a)}{\sum_i L(i) + L(d)} \tag{13}$$


Dietrich, F. (2010). Bayesian group belief. *Social Choice and Welfare, 35*(4), 595-626. doi:10.1007/s00355-010-0453-x

Dietrich, F., & List, C. (2016). 519Probabilistic Opinion Pooling. In A. Hájek & C. Hitchcock (Eds.), *The Oxford Handbook of Probability and Philosophy* (pp. 0): Oxford University Press.

Ulicna, K., Vallardi, G., Charras, G., & Lowe, A. R. (2021). Automated Deep Lineage Tree Analysis Using a Bayesian Single Cell Tracking Approach. *Frontiers in Computer Science, 3*. doi:10.3389/fcomp.2021.734559
